# Evolutionary significance of structural segments trailing beta strand 5 and alpha helix 10 in the design of catalytically active NAL superfamily

**DOI:** 10.1101/505230

**Authors:** P. Manoj Kumar, Prabu Manoharan

**Affiliations:** School of Biotechnology, Madurai Kamaraj University, Madurai, India

## Abstract

The divergence in catalytic actions of N-acetyl neuraminic lyase (NAL) superfamily proteins, all of which have pyruvate as a substrate, suggests common ancestry. Lack of catalytic triad residues essential for binding pyruvate in annotated DHDPS proteins from Gram+ve *B.clausii* (PDBid-3E96) and *O.hiyensis* (PDBid-3D0C) indicated that these proteins are inactive and therefore could be possible early ancestors. Analysis revealed that the most appropriate cavity of these proteins is voluminous and has an elongated topography compared to the trimmed side-wise tilted cavity topography in all other NAL superfamily proteins. Strength and the morphology of the interface regions contouring the cavity are the significant determinants of the topography. It is possible that evolutionary forces led to modulation of the structural segments following the beta strand 5 and alpha helix 10, which are significant participants of interfacial regions. Major structural motions captured by molecular dynamics simulation differentiated the motion of the structural segment following the beta strand 5 of primitive forms as towards the periphery regions of the proteins compared to motion towards core in evolved active forms. We suggest that the motion shift towards the core consequently opened entry channel for substrates and evolution of side-wise tilted cavity.

## Introduction

Proteins have a remarkable property to evolve and acquire new functions. Many modern-day proteins have arisen from a few common ancestors that have evolved continuously over millions of years. Gene duplication and Horizontal Gene Transfer (HGT) are two main avenues that affect evolution at molecular level**^1, 2, 3^** The (α/β) 8 TIM barrel fold is one such domain that evolved through twofold gene duplication from an (α/β) 4 half barrel ancestor with a fused half barrel intermediate**^4,5^**. Copley et al., and Nagano et al., had shown that TIM barrel proteins arisen by divergent evolution from a common stable fold that evolved to show different catalytic activity**^6,7^**. For example, aldolase Type I class of enzymes is a result of one such diversification.

N-acetyl neuraminic lyase superfamily belongs to aldolase Type I class of enzymes. There are six subfamilies in NAL superfamily that include N-acetyl neuraminic acid lyase (NAL), dihydrodipicolinate synthase (DHDPS), 2-keto-3-deoxy gluconate aldolase (KDGA). Typical NAL fold is an (α/β)_8_ TIM barrel with three additional helices at the C-terminus following the 8^th^ alpha helix**^8,9^** The subfamilies of NAL superfamily possibly arose due to HGT that effected transmission of genes across bacterial and archaeal domains resulting in NAL and DHDPS families in bacteria and KDGA in archaea. There have also been instances where a protein like YagE, which is a homolog of KDGA, transferred horizontally to *E.coli* K12 through phage infection**^10, 11, and 12^**. With the common scaffold performing different functions, NAL superfamily is a perfect candidate to study the evolutionary mechanism that resulted in such a divergence.

The underlying unity among these subfamilies is that they bind to common substrate pyruvate, via, Schiff base between the carbonyl oxygen of the pyruvate and the NZ atom of the conserved lysine at the end of the beta strand 6**^13, 14^**. A catalytic triad comprising a strictly conserved tyrosine, a serine/threonine in a strictly conserved GS/TTGE motif and another strictly conserved tyrosine from the neighboring monomer assists pyruvate binding**^15^** Thus NAL superfamily is functional only as biological dimers. The Schiff base forming lysine along with the catalytic triad constitutes primary catalytic residues**^15,16^** Another set of amino acid residues that include a GXD/E motif classified as secondary residues that are loosely conserved, interacts with subfamily specific substrates thus explaining the divergence of NAL superfamily**^17^**. It is likely that NAL superfamily proteins could have evolved from a common ancestor that can bind pyruvate that later diverged to perform varied enzymatic activities. The genetic material of such an ancestral form might have probably transmitted to other domains of life through HGT with mutations in due course of time enabling divergence into various subfamilies**^17^**

Protein-protein interaction gives rise to oligomeric assemblies for mediating various cellular functions. Interfacial region constitutes the burial of the solvent-exposed hydrophobic/hydrophilic surfaces of the partaking residues from both the subunits. Chen et al. 2013 had shown a direct correlation between binding affinity between subunits and increased buried surface area at the interface**^18^**. Thus buried surface area is indicative of the strength of the interface which in turn indicates stable assemblies. Effective burial of the surfaces depends on two primary considerations: geometrical factors such as interface size, planarity, sphericity and complementarity and chemical nature of the participating amino acids such as hydrophobicity, electrostatics, and hydrogen bonding**^19^**. Hydrophobic effect plays a determinant role in the formation of an interface, although the effect is less pronounced than seen in the protein folding**^20,21,22,23^** Polar residues also play a significant role in the formation of interface contributing through hydrogen bonding and salt bridges**^23,24^**. This tight packing of the buried surfaces is due to a significant number of partaking interfacing residues, a high degree of hydrophobicity and active stabilization of polar groups through hydrogen bonding and salt bridges. However, the increased degree of polarity results in a poor packing of atoms thereby weakens the interface of the proteins**^25^**. This observation is evident from the fact that homodimers are more closely packed compared to loosely packed heterodimers, as hydrophobic residues dominate the former while the latter by hydrophilic residues**^26^**. The more significant number of interfacing residues linearly increases the interface area. Larger interface area generally results in stronger protein assemblies**^27^**. Burial of fewer atoms results in weak interfaces characterized by poor packing and increased solvent accessibility**^28^**. Solvent accessibility is one of the significant determinants of local protein flexibility and dynamics**^29^**. Protein complexes with a relatively larger solvent accessible surface area in their interface will be weakly associated. Such complexes will be more dynamic and exhibit conformational fluctuations among the partaking subunits.

Shape complementarity between the subunits is a key factor that leads to stronger oligomeric assemblies**^30, 31, and 32^**. Packing of atoms to form a stronger interface results when the residues from both the monomers either protrude from the surface exhibiting convexity in their surfaces**^26,33,34^** or exhibiting shape complementarity between surface concavity in one subunit and the surface convexity of the interacting partner**^35^**. Poor internal packing results when there is no shape complementarity between the partaking surfaces giving rise to cavity pocket in between the interface region**^36, 37, 38^** Cavities as they result from defects in packing would have a profound negative effect in the buried surface area and free energy of folding and binding, thereby destabilizing the structures**^39^**. Cavity filling and the cavity forming mutagenesis experiment had already established the negative correlation between the formation of cavities and stability of fold and assemblies**^39, 40, 41, 42^**. Though there is an energetic cost, proteins tolerate the formation of cavities because of compelling functional necessities. Stronger subunit interactions though stabilize the structure, may be deleterious to the proper functioning of the proteins. Therefore formation of a cavity is often a stability-function tradeoff. Lee et al., 1999 had demonstrated that though cavity filling mutations contribute to the conformational stability in α1AT, heavily hamper the serine protease inhibition activity**^41^**. Presence of cavities as they weaken the interface often leads to rearrangement of the subunits that may facilitate the binding of ligands or may generate channels to transport the ligands to and from the active site of proteins**^43, 44^**

Protein dynamics can provide a cassette of conformational ensembles that populate the free energy landscapes**^45, 46^** enabling the protein to have multiple functional states. The most populous conformation, which is usually also the most stable, determines the native function of the protein. Evolutionary changes can shift the population from one pre-existing most populous conformation to another, thereby leading to the emergence of novel enzymes. Thus Conformational dynamics is the fulcrum of divergent evolution towards producing new enzymes**^47, 48^**. It exhibits through any one of the processes ranging from minor fluctuations such as assuming different side chain rotamers, flexibility in active site cavity, and movement of secondary structure elements to even major rearrangement of tertiary structural domains and quaternary subunit assemblies**^46^**. Structural determinants such as disorders, low compactness, faults in the local folding due to poor tertiary interactions solvent accessibility and loose packing in the interface are the main factors for conformational dynamics**^25^**. Franzosa and Xia, 2009 had correlated the constraints imposed upon by these structural factors on the residual evolution rate**^49^**. Solvent accessible residues have possibly weaker evolutionary constraints and therefore are more prone to mutational changes compared to buried residues**^49^**. The mutational effect on function and stability is a crucial factor in understanding the evolutionary dynamics of proteins. Neutral mutations effects non-adaptive changes on protein function and stability and therefore are generally not destabilizing**^50^**. The tolerance limit of the fold towards mutations is proportional to the robustness of the proteins**^51^**. The mutational robustness is measured by a fitness landscape that depicts the epistasis effect; wherein deleterious mutations inflict negative epistatis effect while neutral mutations reinforce the positive effect. Thus neutral mutations buffer the negative epistatis to keep the fitness landscape intact thereby defining the robustness**^52,53^**. Chan et al., 2017 had demonstrated the robustness of the TIM barrel fold whose sequence and structural constraints defines the correlation between the fitness landscapes of distant orthologs**^54^**. Conformational plasticity in the active site is the major determinant of change which results in the evolutionary divergence. Thus the inducing of functional divergence of proteins without affecting the stability of the scaffold is a stability-function tradeoff**^50^**. Toth-Petroczy and Tawfik, 2014 defines a term called ‘Polarity’ as a key to innovability**^51^**. It measures the degree of how well the active site and stability defining scaffold are separated and autonomous. Therefore higher the ‘Polarity’ more will be the divergence of a new function by the same fold. In robust folds such as TIM barrel, the well separated rigid high order scaffold and the flexible active site **^55, 56^** facilitates the mutation in the active site while not affecting the scaffold stability. The mutated active site under a stable scaffold forms the basis for the divergence and innovability of NAL superfamily proteins. Mutations that operate through deletions and insertions of residues, driven by solvent accessibility and low packing density, may ruin or weaken the existing interface and create new stronger interfaces. Thus these operations are mechanistic factors of evolution that lead to oligomerization or even to the emergence of novel proteins**^57^**.

Here we report a possible ancestral form of protein with NAL scaffold that is seemingly inactive and the likely evolutionary development of active NAL superfamily proteins with divergent functions. The insight gained on the cavity topography, and interfacial regions can provide a knowledge resource in the use of directed evolution for developing novel NAL superfamily proteins. Horowitz 1945 and Jensen 1976 had proposed that enzymes within similar metabolic pathways should be homologues**^58, 59^**. The participation of different NAL superfamily proteins which have similar catalysis mechanism in various metabolic pathways suggests that understanding the evolution of these proteins may shed light on metabolic pathway evolution.

## Materials and Methods

### 1. Naming and Definition

Selected NAL superfamily proteins from DHDPS, NAL and KDGA subfamilies reported in PDB *(https://www.rcsb.org/)* are analyzed (The proteins are listed in table ST1 in supplementary information SI1). Only the results of the analysis of proteins 3E96, 3D0C, and 3B4U (all annotated DHDPS), 1O5K (DHDPS), 1F5Z (NAL), 1W37 (KDGA) is presented. Since a tyrosine from neighboring monomer takes part in the catalytic mechanism, the biological dimer of the proteins was analyzed. Monomer labeled A is colored yellow and monomer labeled B is colored red throughout the article. The secondary structural segment that follows the beta strand 5 is named depending on whether it is loop or helix, as loop A/helix A of monomer A and loop B/helix B of monomer B. The loop segment that trails the helix A and B are also the area of interest. The secondary structural segment of monomer A and B that follows the beta strand 4 is named as loop A1 and loop B1, respectively. The tyrosine from the neighboring monomer that takes part in catalytic triad is harbored on this segment. The secondary structural segment that follows the alpha helix 10 is named as loop A2/helix A and loop B2/helix B. (figure SF1 of supplementary information SI1 illustrates the naming of the segments)

Since all the proteins are homomers, the physiochemical properties will be similar in both the subunits. Therefore only the active site cavity of monomer A is discussed. The secondary structural segments loop A/helix A, loop A2/helix A2, and loop B/helix B and loop B1 that are part of the active site of monomer A is considered. Segment loop B/helix B is also taken for analysis because this segment has a significant role to play as will be discussed subsequently.

### Multiple sequence alignment

Structure-guided multiple sequence alignment was done to check for the conservation of primary and secondary catalytic residues and the omissions and additions in the segment trailing the beta strand 5 and alpha helix 10. Selected PDB entries were extracted as monomers and superimposed in Chimera60 (Petterson et al., 2004; https://www.cgl.ucsf.edu/chimera/), followed by structure-guided multiple sequence alignment.

### 3. Active site cavity volume and topography analysis

NAL superfamily proteins possessing same fold but divergent in functions put forth a question whether active site cavity volume and topography is the deciding factor for the divergence. Active site cavity volume calculation and topography mapping of the selected PDB structures of NAL superfamily was done using CastP**^61^** (Computed Atlas of Surface Topography of Proteins). Solvent molecules including water molecules are removed before calculations. CastP lists all the cavities present in the protein with details of volume, area and the residues present in the cavity. The most appropriate cavity or the active site cavity is the one that lists almost all catalytic residues. Probe radius was set at 2.0Å for calculation as because only at this radius, the most appropriate cavity of the proteins could be identified.

### 4. Solvent accessible surface area analysis (SASA) on active site cavity contouring interfacial regions of the proteins

The residues from loop A/helix A, loop B/helix B, loop B1, loop A2/helix A2, loop A1, and loop B2/helix B2 was listed as separate set PDB coordinate file I. Another set PDB coordinate file II lists the residues from the same segments excluding the residues from loop A2/helix A2 and loop B2/helix B2. Generation of separate files was done to identify the actual contribution of these segments to active site cavity contouring interfacial regions since the selected segments also interact with other insignificant regions. Solvent accessibility and buried surface area due to interface formation was determined using PDBePISA server (http://www.ebi.ac.uk/pdbe/pisa/). The difference in the values between the two coordinate file sets is computed to know the contribution of each segment towards interface formation. The accessible and buried surface of the segments of only one monomer was presented and discussed as the monomers of a homodimeric protein exhibit identical behavior in their physicochemical properties. Solvation free energy gain due to interface formation (Δ^i^G) which reflects the tightness of interface was also computed using PDBePISA. Negative free energy values expressed in Kcal/Mol indicate hydrophobic interface or positive protein affinity. In other words, larger the negative free energy value, the better the local folding and hence tighter the interface.

### 5. Solvent Accessible Surface Area (SASA) analysis on segment following beta strand 5

Solvent accessible surface area (SASA) and Buried surface area (BSA) of the residues harbored on the segment loop A/helix A (loop A/helix B) that followed the beta strand 5 were computed using AreaIMOL of CCP4. Only the segment in monomer A (loop A/helix A) is discussed to avoid repetition. SASA calculations for both dimeric and monomeric form of proteins were determined. The ASA value of a residue in the dimeric form is the total area, and the value in monomeric form is the actual ASA. The difference between the SASA values of the residue as the dimer and as the monomer is the BSA value of that residue. The percentage accessibility and the buried surface of each residue of the segment was determined (Supplementary information SI2: SF2, ST2)

### 6. All-Atom Molecular Dynamics Simulation

The MD simulation studies on Biological dimers of proteins 3E96, 1W37, 2V8Z (YagE), and 1NAL (NAL) was done. The 50ns simulation work reported in this study is performed using Gromacs (Version 4.6.6)**^62^** with the gromos54a7**^63^** force field. The crystallographic water present in the structures removed and hydrogen atom added. The system was solvated using the simple point charge (SPC) **^64^** water model in a cubic cell with a minimum distance of 12 Å between the solute and each face of the box. The system was neutralized by adding NaCl with an ionic concentration of 0.15 M. At this stage, the simulation model constructed for 3E96, 1W37, NAL, and YagE comprises one homodimer of each. The system was energy minimized for 2000 steps initially by steepest descent followed by the conjugate gradient. The NVT equilibration performed for a 100 ps, by applying the velocity-rescaling thermostat**^65^** the protein backbone restrained and the solvent molecules are allowed to equilibrate around the protein. Following the equilibration to a target temperature, 1 ns of NPT equilibration performed, by applying the Nose-Hoover thermostat**^66, 67^**. The Parrinello-Rahman barostat**^68^** was used to couple pressure isotropically, to a value of 1.0 bar. The simulation was performed at constant pressure and temperature (300 K) with a time step of 2 fs. The final snapshot of equilibration run was used to start the 50 ns NPT production run with position restrain on protein backbone removed. The non bonded interactions were truncated at 8 Å cut off and the neighbor list updated every 1 step. Long-range electrostatic interactions were calculated using particle mesh Ewald (PME)**^69^** method. All bonds are constrained using the linear constraint solver (LINCS)**^70^** algorithm.

Essential Dynamics**^71^** (EM) was performed using Principle Component Analysis (PCA) a multivariate technique was used to capture the domain motions that are fundamental to the protein function. Only the backbone atoms were included during the EM study. The extreme motions captured with the help of EM were used to identify dynamic domains. Major protein motion that contributes to the overall motion was visualized using modevector script in pymol.

## Results

### Proteins 3E96 and 3D0C should be non-functional

Multiple sequences alignment showed that the proteins 3E96 and 3D0C annotated as DHDPS enzymes lack essential catalytic residues (Figure 1). The strictly conserved lysine harbored at the end of the beta strand 6 that forms Schiff base with carbonyl carbon of pyruvate is conserved. However, the catalytic triad necessary to abstract and shuttle the proton from the carboxyl group of the pyruvate to and from the solvent is not present in these proteins. One of the triad residues, the strictly essential tyrosine is conserved in proteins 3E96 and 3D0C. However, the other two essential triad residues threonine or serine in the GS/TTGE signature motif and tyrosine from the neighboring subunit are not conserved in these proteins. Notably, there is a histidine residue in the place of the tyrosine. Unlike the tyrosine, which is mapped in the disallowed region of the Ramachandran plot and whose side chain is oriented and reaches into the active site of the neighboring subunit, the histidine, whose dihedral angles falls in the allowed region of the Ramachandran plot, orient its side chain towards its own subunit (Figure 2). In all active proteins, the catalytic triad residues are arranged like a triangle, with the hydroxyl group of each residue forming the vertices (Figure 2). The absence of triad residues and its triangle geometry arrangement thus makes proteins 3E96 and 3D0C unviable for binding pyruvate. Proteins 3E96 and 3D0C also lack the GXD motif, necessary to bind the subfamily specific secondary substrate. Most importantly MSA also revealed that the subfamily specific residue viz., Arginine in DHDPS, Leucine in NAL and Alanine in KDGA is not present in these proteins. Thus these proteins cannot be classified under any of the subfamilies and therefore should be nonfunctional for NAL superfamily catalysis.

**Figure 1:**
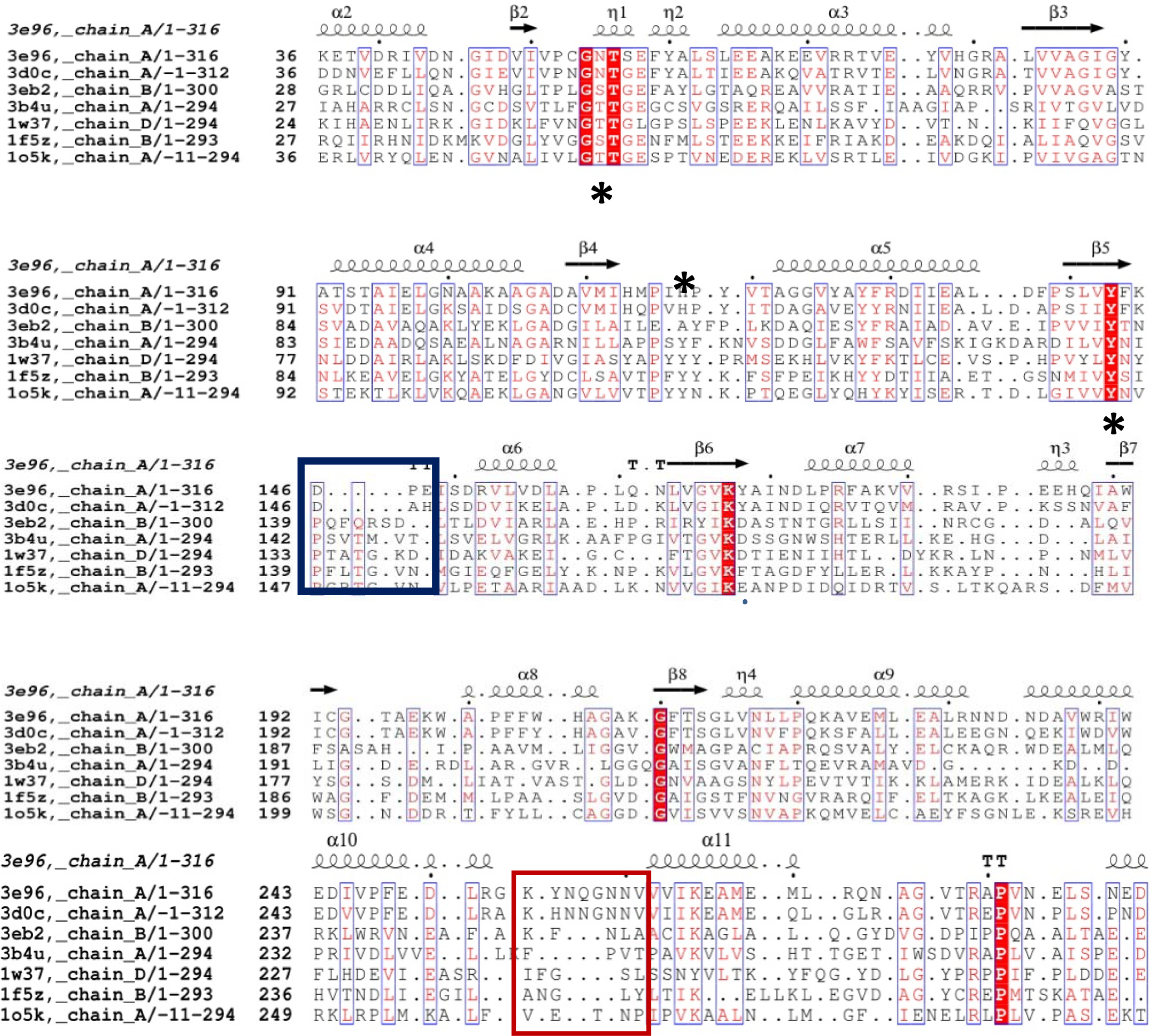
Structure based multiple sequence alignment showing the unique features of protein 3E96 and 3D0C. Residues boxed in black are primary catalytic residues involved in binding primary substrate pyruvate and the catalytic triad residues are marked with asterisk symbol. Except tyrosine in beta strand 5, proteins 3E96 and 3D0C lack triad residues. GXD motif in the dark blue block is involved in binding subfamily specific secondary substrates, which is absent in proteins 3E96, 3D0C, 3B4U and 3EB2. Loop A/helix A (loop B/helix B) segment that trail the beta strand 5 is boxed in green. Proteins 3E96 and 3D0C have relatively more deletions compared to rest of the proteins. Evolution hot spot residues Aspartate (Asp146) and Glutamate (Glu148) is present in these proteins but absent in rest of the proteins. Loop A2/helix A2 (loop B2/helix B2) segment that trail the alpha helix 10 is boxed in light blue. There are relatively more additions in proteins 3E96 and 3D0C, compared to rest of the proteins

**Figure 2:**
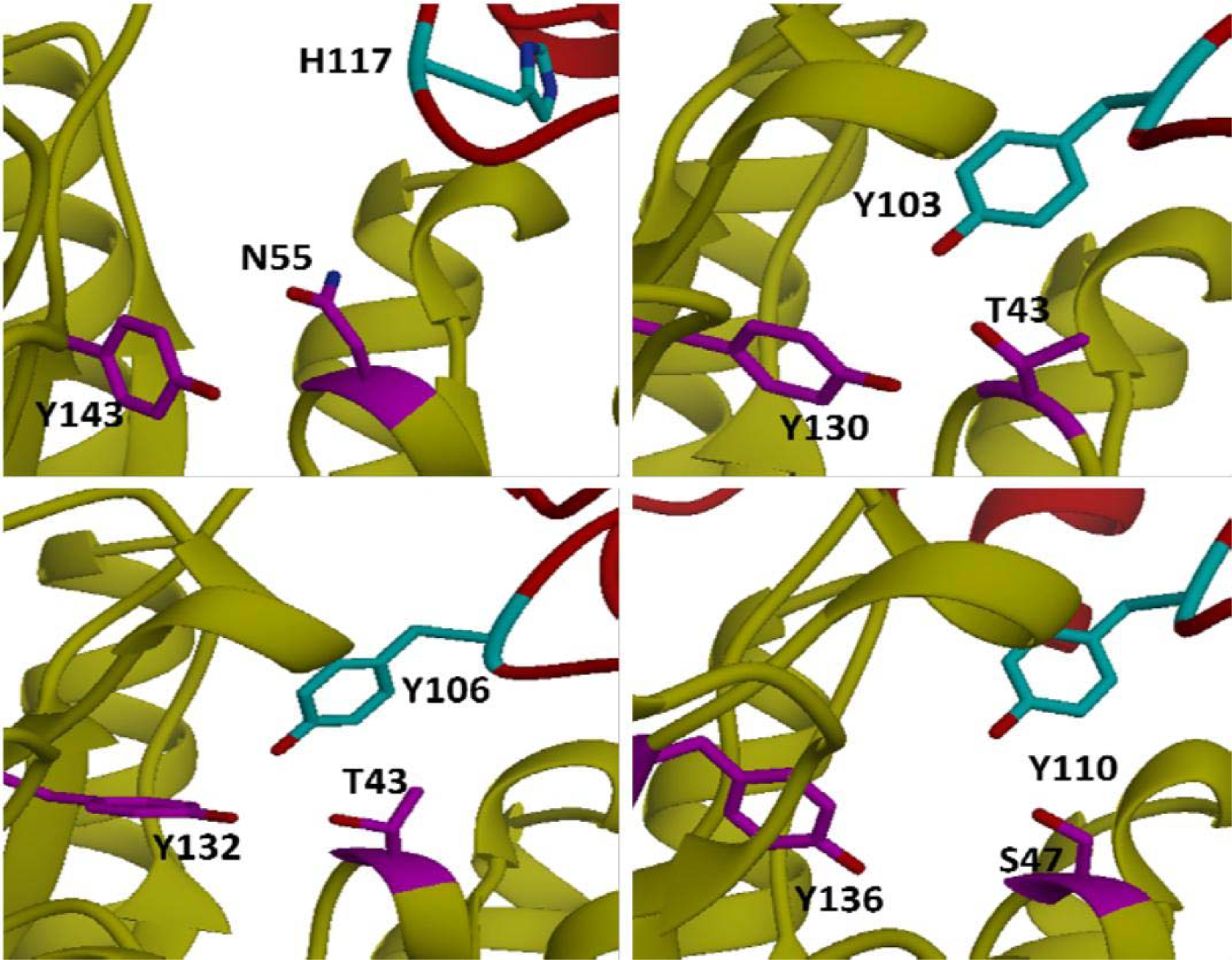
Catalytic triad arrangement formed by Serine/Threonine, Tyrosine and another tyrosine from neighboring monomer. Top left (3E96), Top right (1O5K), Bottom left (1W37); Bottom right (1NAL). Protein 3E96 possessing a histidine (H117) which is oriented towards its own monomer is shown, indicating the absence of catalytic triad

### Partially functioning proteins with pyruvate binding ability

Multiple sequence analysis revealed that annotated DHDPS proteins (PDBid-3B4U and 3EB2) have the conserved primary catalytic residues necessary for pyruvate binding such as Schiff base forming lysine, and catalytic triad residues with appropriate triangle topology. These features render these proteins to bind pyruvate. Protein 3EB2 lack the GXD motif, while protein 3B4U possess a GDE motif, and seems well qualified to mimic DHDPS activity. However, these proteins lack the essential Arginine needed for binding L-ASA**^72^**. Thus these proteins are partially functional to bind pyruvate but cannot bind the subfamily specific substrates.

### Extensive deletion in beta strand 5 and insertion in alpha helix 10 in proteins 3E96 and 3D0C

Multiple sequence alignment revealed that there is an extensive deletion in loop B (loop A), the segment that trails the beta strand 5 in proteins 3E96 and 3D0C compared to relatively large number of residues in the counterpart segment loop B/helix B (loop A/helix A) in rest of the proteins (Figure 1). The signature residue, viz., Arginine in DHDPS, Alanine in KDGA and Leucine in NAL are present in this segment. Residue in alignment with this signature residue is vacant or deleted in proteins 3E96 and 3D0C (Figure 1). Notably, there is an aspartate and a Glutamate (it is histidine in 3D0C) in this segment in proteins 3E96 and 3D0C which is distinctly conspicuous and are absent in rest of the proteins. In contrast there is an extensive deletion in loop A2/helix A2 (loop B2/helix B2), the segment that trails the alpha helix 10, in rest of the proteins compared to the relatively large number of residues in the counterpart segment, loop A2 (loop B2) in proteins 3E96 and 3D0C (figure 1).

### Loose interface segment 1 (IS1) and tight interface segment 2 (IS2) in protein 3E96 and 3DOC compared to tight IS1 and loose IS2 in rest of the protein

Analysis on the biological dimers made using PDBePISA server showed that dimeric interfaces are unique in the proteins 3E96, which is open in the core and closed in the periphery, compared to closed core and opened periphery in rest of the proteins, including the partially active protein 3B4U (Figure 3; Image A in all the panels). The interfacial segment in the core, henceforth called as IS1, is formed by residues from left ends of loop B1 of monomer B with loop A/helix A of monomer A (Figure 3; Image B in all the panels). The triad partaking tyrosine harbored on left end of loop B1 and a subfamily specific residue, alanine of KDGA, leucine of NAL, and Arginine of DHDPS, harbored on loop A/helix A are interface partners. In DHDPS subfamily (1O5K), the residues from loop B/helix B also contributes to IS1 strengthening it further compared to proteins of KDGA and NAL subfamilies (Figure 3; image A in panel V). Though the loopB/helix B segments in proteins of NAL and KDGA subfamilies do not contribute, the segments are close to loop A/helix A, making the IS1 region steric hindered. In proteins 3E96, the segments are far apart and therefore do not form interface except for a small surface contact between a Lys145 and a Pro118 residue harbored on loop A and loop B1 respectively, hence there is none or significantly small IS1 in these proteins. The interfacial segment in the peripheral region named as IS2 is formed by burial of surfaces of residues from loop A2/helix A2 and the right end of loop B1 (Figure 3; Image C in all the panels). In proteins 3E96 and 3D0C, in addition to residues from loop A2 and loop B1, residues Asp146 to Glu148 from the right end of loop B also take part in IS2. In rest of the proteins, IS2 is negligibly smaller except for a small surface contact between the triad partaking tyrosine from loop B1 and another aligned residue () in helix A2 (Image C in panel III). In some proteins such as active DHDPS from *C.glutamicum*, the segments are far apart that there is no formation of IS2 as indicated by the absence of surface contact between the segments (image not shown). An important observation that may have evolutionary significance is the surface morphology of IS2, which is a shallow concave trough (Figure 4). In protein 3E96, the partaking residues from loop B, Asp146 to Glu148, are projected downwards thereby demonstrating a trough orientation (Panel II, Figure 4). Also, the orientation of the loop A2 which is protruding towards the trough topology segment of loop B (Panel II, Figure 4) resulted in the reduced distance that led to the burial of their surfaces towards the formation of IS2 (Panel III, Figure 4).

**Figure 3:**
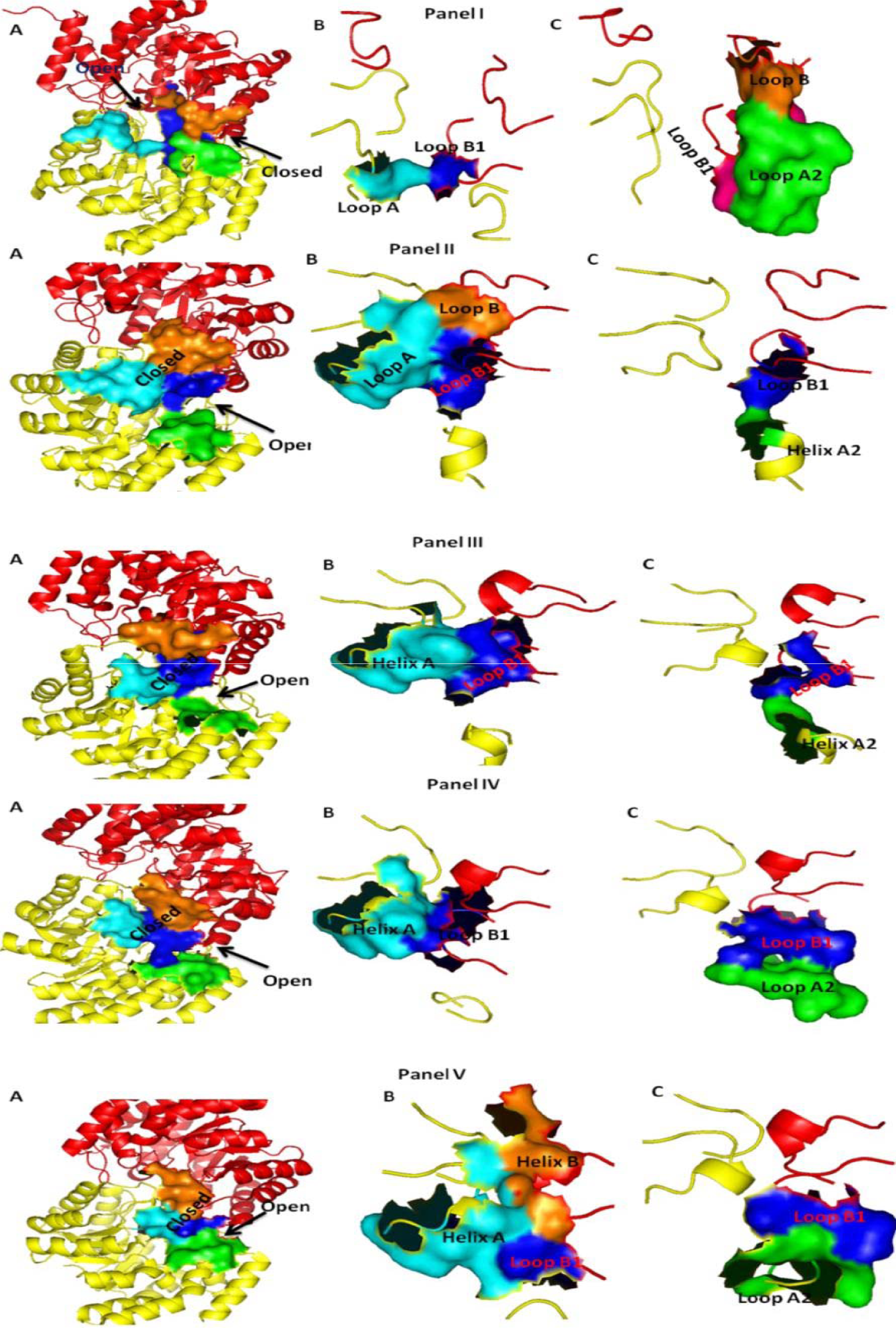
Panel I (3E96), Panel II (3B4U); Panel III (1F5Z); Panel IV (1W37); Panel V (1O5K). Image A in all the panels shows the interface scenario in each protein. Protein 3E96 has open core and closed periphery and vice-versa in rest of the proteins. Image B in all the panel shows the IS1 region. Panel I shows a small interface in protein 3E96, all other proteins have large IS1 region as shown in panel II to V. Panel II and V shows that helix B in proteins 3B4U and 1O5K also contribute to the ISI interface. Image C in all the panel shows the IS2 region. Panel I (3E96) shows a large IS2 region compared to rest of the proteins as depicted in panel II to V. As indicated in Image A in panel III and IV, the helix B (Orange coloured surface) though not taking part in the interface IS1, provides steric hindrance in the core in proteins 1F5Z (NAL) and 1W37 (KDGA)

**Figure 4:**
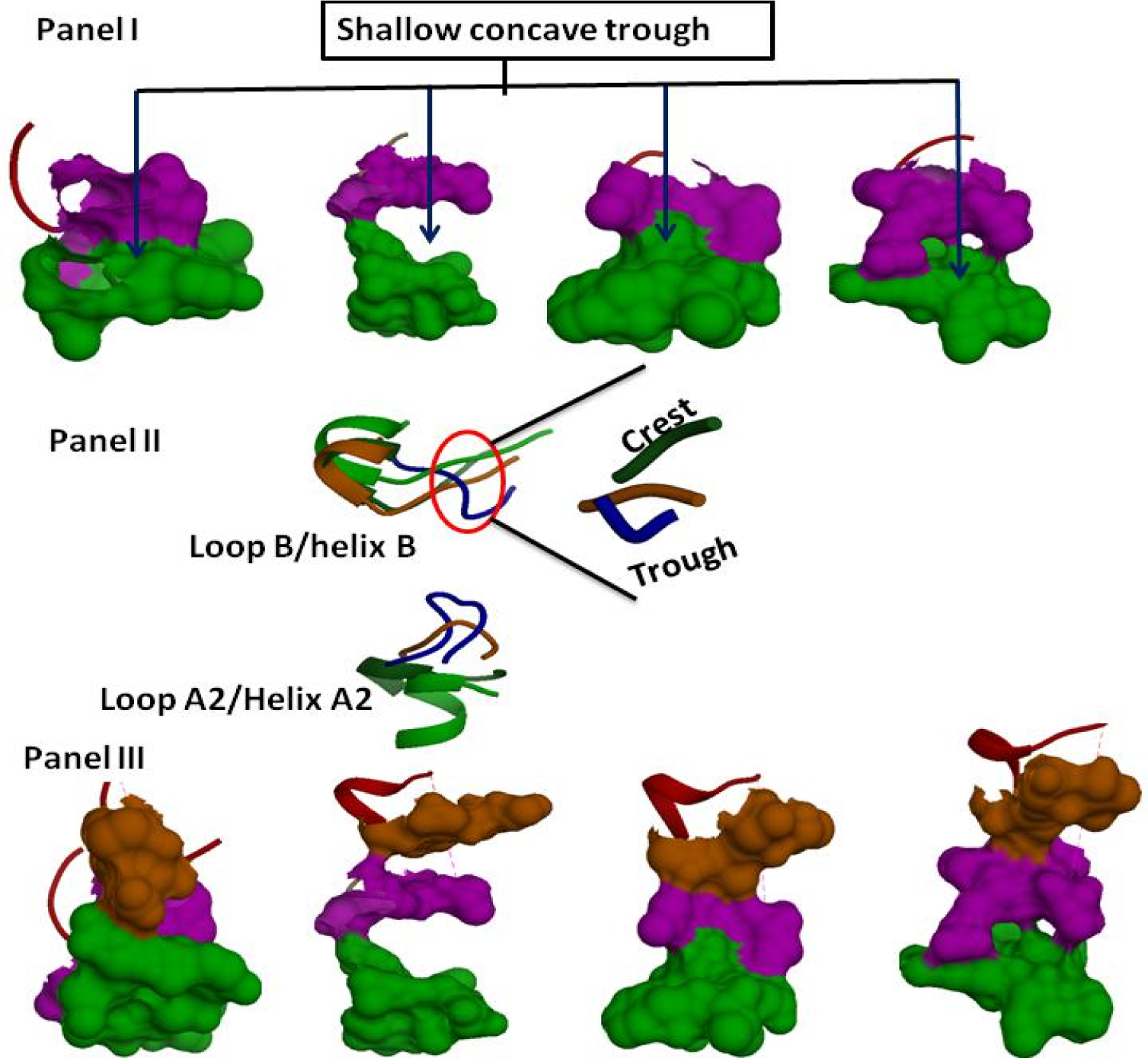
Panel I showing the morphology of the surface contact of loop A2/helix A2 (Green) and loop B1 (Magenta) forming a shallow concave trough. (from left to right) 3E96, 1F5Z, 1O5K, 1W37. Panel II shows the loop segment trailing the loop B (residues D146-E148) in protein 3E96 (blue) taking a trough orientation, while the corresponding structurally aligned segments in other proteins takes a crest topology (Green-1W37; Orange-1F5Z). Panel III showing the surface of trough orienting segment of loop B in protein 3E96 (Orange) make contact with loop A2 (green), rendering the shallow concave trough sterically hindered (first from the left). While in rest of the proteins (second from left-same order as is given in panel I) the surfaces of the crest oriented segment of the loop trailing helix B (orange) do not make surface contact with loop A2/helix A2 (green) and therefore does not encroach the shallow concave trough.

The segment in the rest of the proteins that are in alignment with the trough orienting segment of loop B of protein 3E96 is projected upwards demonstrating crest orientation (Panel II, Figure 4). Unlike the elongated and upward orientation of loop A2 in protein 3E96 and 3D0C, the loopA2/helix A2 in rest of the proteins is flattened (Panel II, Figure 4). Crest topology of loop segment that trail the helix B and relatively flattened loop A2/helix A2 results in the relatively large distance between the segments and therefore their surfaces do not involve in interface formation and therefore are not constituents of IS2 (Panel III, Figure 4).

Weaker IS1 and stronger IS2 in protein 3E96 and 3D0C and opposing scenario in rest of the proteins is further corroborated by percentage buried surface area (%BSA) and solvation free energy gain due to interface formation (ΔG, Kcal/mol) determined by PDBePISA server (**supplementary information SI3**: SF3, ST3)

### Unique Cavity topography in proteins 3E96

Surface topography mapping of the active site cavity of the selected proteins done using CastP showed that protein 3E96 and 3D0C has a large cavity with an elongated vertical topography (Figure 5). The identified cavity in protein 3E96 is not restricted within the monomer; say A but spread across the monomer B also (Figure 5). CastP calculations showed that the identified cavity in monomer A of protein 3E96 has the larger cavity volume of about 1820Å^3^compared to the cavity in monomer B is just 470.8Å^3^ This is because the cavity of monomer A transcends across monomer B and therefore the cavity in monomer B is restricted from spreading further resulting in reduced cavity volume (Figure 5). Thus there is no distinct cavity in each monomer. The surface contact between Lys145 and Pro118 that forms the IS1, contour the cavity in depth in the core and the surfaces of IS2 contour the cavity in the peripheral region of the protein (Figure 5). Active site cavity in rest of the proteins, including the partially functioning proteins, irrespective of their affiliations, is trimmed in volume and has a side-wise tilted topology (Figure 5). In some DHDPS proteins, the cavity is highly trimmed to have average to least active site volume. Each monomer of the biological dimer has a distinct active site cavity with identical topography and volume (Figure 5). The surface of IS1 contours the cavity, forming the ‘roof,’ in the core, while the surface of IS2 contours the cavity in depth in the peripheral region of the protein. Electrostatic potential surface mapping showed that the IS2 region contouring the cavity is electronegative in proteins 3E96 (Figure 6). The major contributors to the electronegativity of IS2 in protein 3E96 are Asp146 and Glu148 both harbored on the trough oriented segment of loop B. Opposing scenario is seen in IS2 region which is electropositive in rest of the proteins (Figure 6). These features suggest that electronegatively charged pyruvate can access the entry channel of the rest of the proteins, and may be restricted in 3E96. (Selection criteria, the volume of the cavity and the constituting residues are given in the supplementary information SI4: ST4, ST5).

**Figure 5:**
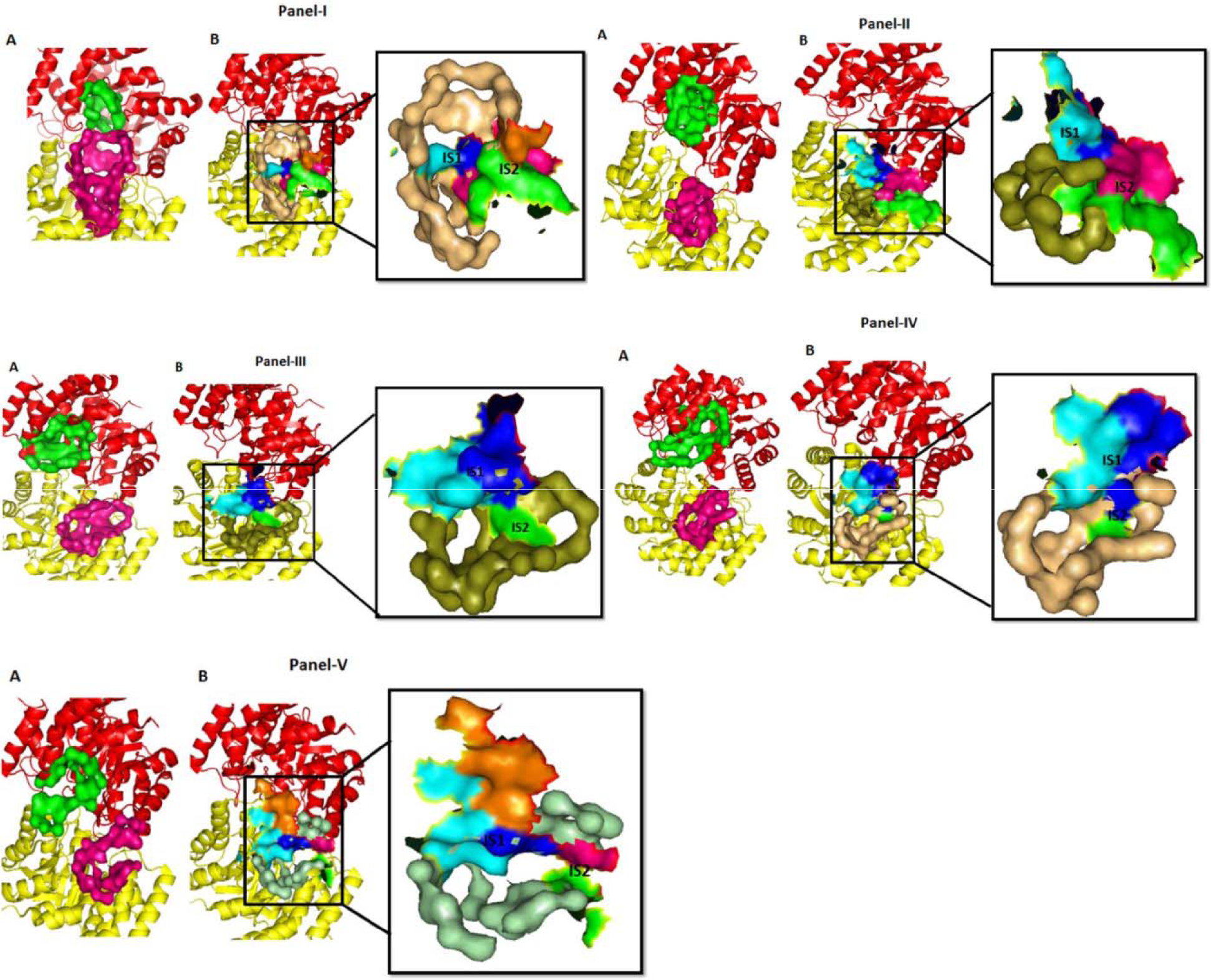
Surface mapping of active site cavity topography in proteins 3E96 (Panel I), 1W37 (KDGA; Panel II), 1F5Z (NAL; Panel III), 3B4U (annotated DHDPS; Panel IV), 1XXX (DHDPS; Panel V). Image A in each panel shows that the proteins possess individual active site cavity for each subunit of the biological dimers, with identical topography, except in protein 3E96. Image B in panel I shows elongated vertical cavity topography in protein 3E96, while side-wise tilted topography in rest of the proteins, including the partially active 3B4U. Panel B also illustrates the surfaces of IS1 and IS2 contouring the cavity. In protein 3E96, the surface of IS1 forms the backbone of the cavity, allowing the cavity to transcend the neighboring subunit. In rest of the proteins, the IS1 region forms the ‘roof’ of the cavity, thereby restricting it from spreading to neighboring subunit, thus giving rise to two distinct active site cavities. The shallow concave trough of IS2 region forms the back bone of the cavity in rest of the proteins and therefore does not restrict the side-wise spreading cavity, while the tight IS2 region in protein 3E96 restricts the cavity expanding side-wise.

**Figure 6:**
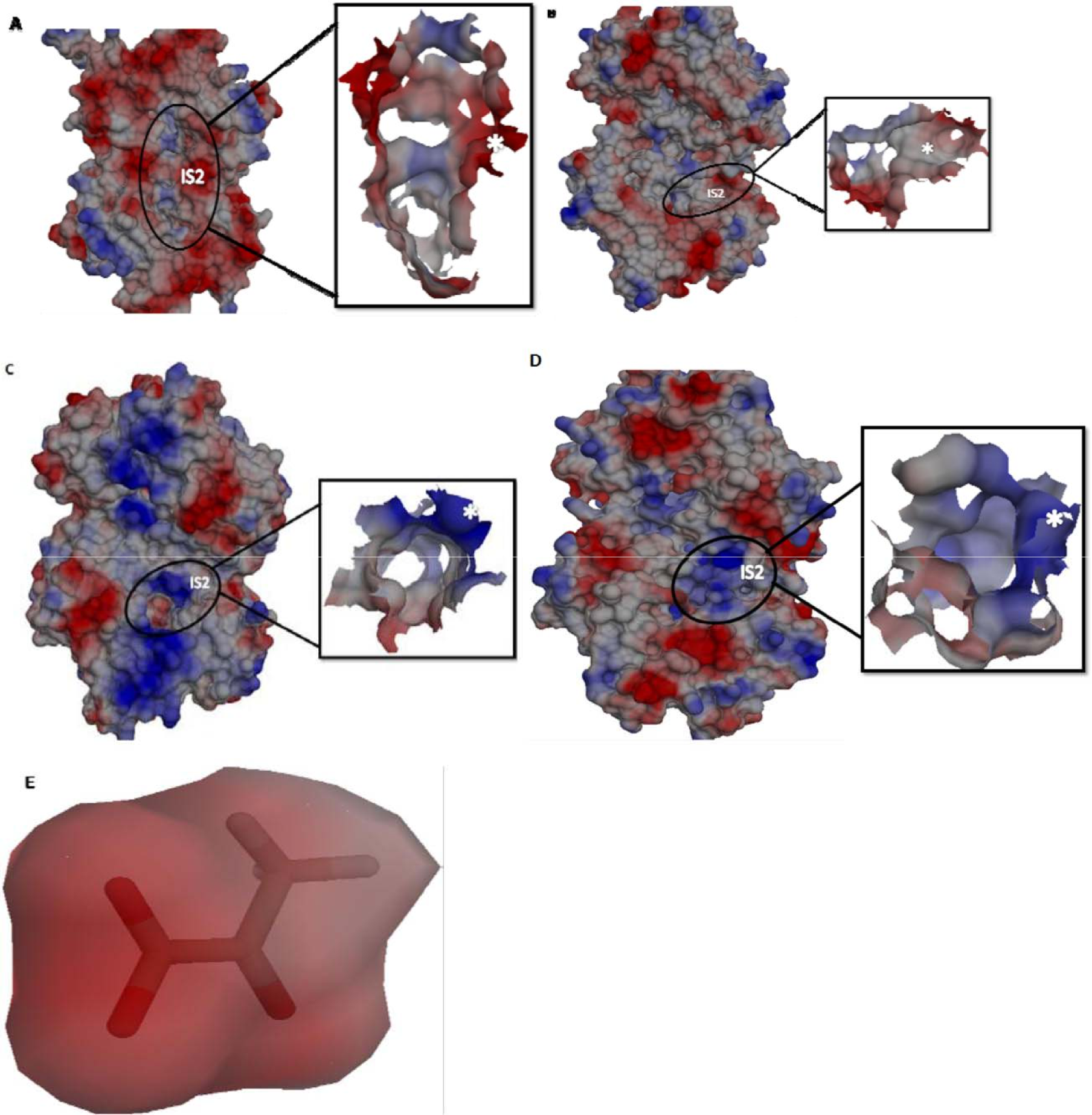
Columbic electrostatic potential mapped on the surface of the proteins (A) 3E96, (B) 1F5Z (NAL), (C) 1O5K (DHDPS) and (D) 1W37 (KDGA). Electronegative potential of IS2 region of protein 3E96 compared to positive electrostatic potential of the IS2 of rest of the proteins can be clearly differentiated. Electrostatic potential of the surface of the cavity is shown in the inset with the potential in IS2 region is indicated in white asterisk mark. (E) Negative electrostatic potential of pyruvate is shown.

### Subfamily-specific residue harbored on loop A/Helix A in rest of the proteins is highly solvent inaccessible

The SASA analysis on the segment that trails beta strand 5, the loop A (loop B) of proteins 3E96 and 3D0C and loopA/HelixA (loop B/helix B) done using AreaIMOL of CCP4 suite revealed that the segment in proteins 3E96 and 3D0C are highly solvent accessible compared to the same segment in rest of the proteins. More than 80% of the surface area of this segment in these proteins is solvent accessible (Figure 7A). In partially functioning proteins such as 3EB2 and 3B4U, 70% of the surface area of this segment is solvent exposed (Figure 7A). The segment in NAL subfamily proteins has their 80% of the surface exposed to the solvent which is closely followed by proteins of KDGA subfamily. In DHDPS proteins the segment is relatively mostly solvent inaccessible as evident from the least percentage of this segment area accessible for solvent (Figure 7A). (Numerical values and chart showing percentage ASA and BSA for rest of the proteins including 3D0C and 3EB2 are given in supplementary information SI5: SF4, ST6-ST11). The signature residue that led to subfamily classification, viz., alanine of KDGA and leucine of NAL which are the most highly buried residues, among all the residues in that segment. However in DHDPS though the signature residue arginine is profoundly buried, the strictly conserved threonine which is next to arginine is the most highly buried residue. Remarkably the KDGA and NAL proteins too harbor a serine or threonine prominently, next to their respective signature residue, which is also primarily buried. The percentage area of the buried surface area of this subfamily specific residue is large compared to their solvent accessible area. There prevails a trend wherein the percentage of BSA increases and reaches the maximum for the signature specific residue (in DHDPS the maximum is for the next residue threonine) and decrease to become completely accessible. This phenomenon is not seen in protein 3E96. When ignoring the gaps in MSA, the residue in alignment with the subfamily specific residue is glutamate in 3E96. SASA analysis showed that this residue is mostly solvent accessible. However, the percentage area of buried surfaces of the residues is relatively large compared to the other residues harbored on the segment in these two proteins. Percentage ASA and BSA of individual residues harbored on the segment in selected proteins are given in figure 7 (Numerical values are given in supplementary information SI5).

**Figure 7:**
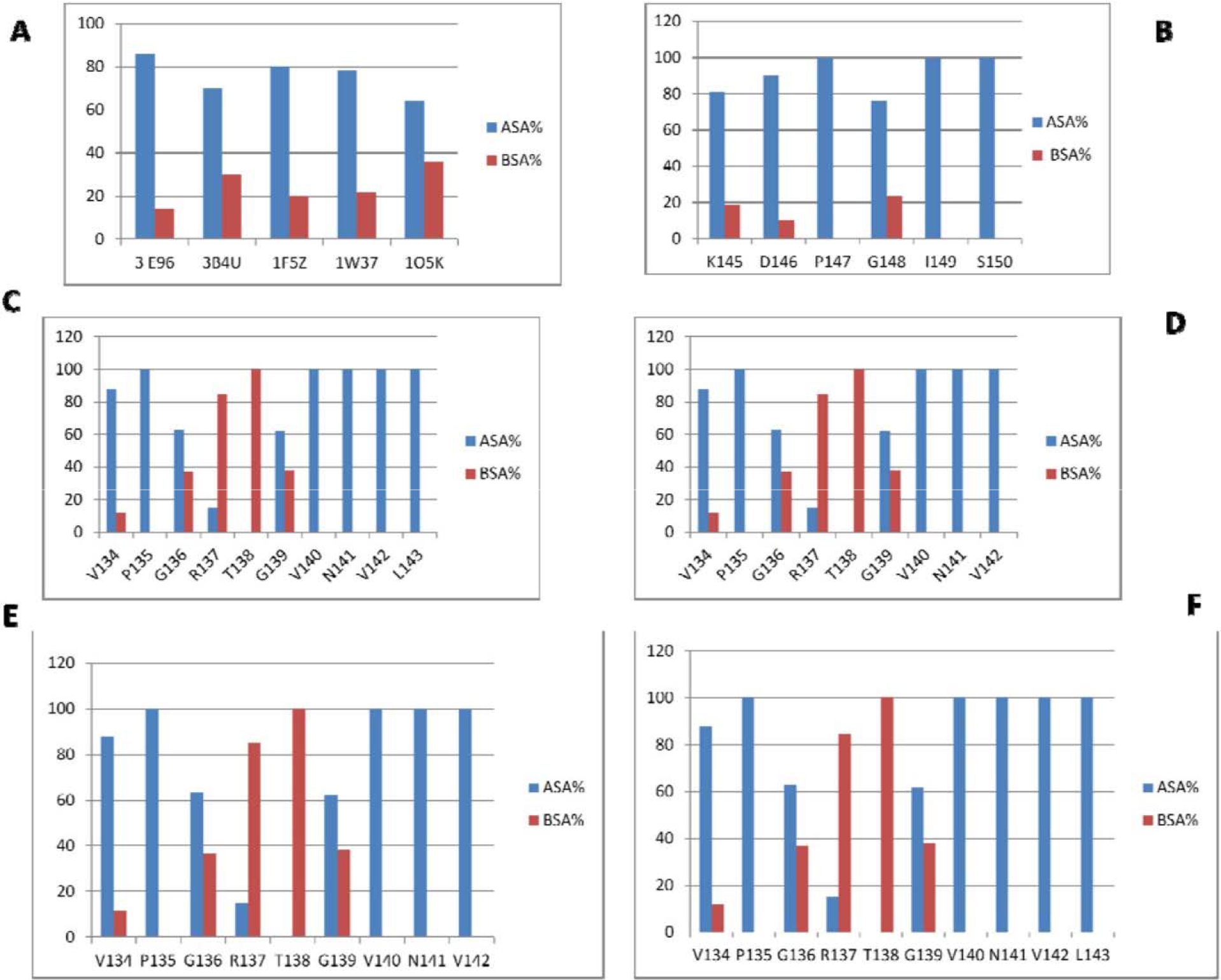
(A) Percentage of accessible surface area (ASA) and buried surface area (BSA) of the beta strand 5 (loopA/helix A) in selected proteins of NAL superfamily. Percentage ASA and BSA of individual residues harboured in the beta strand 5 of proteins 3E96 (B), 3B4U (C), 1F5Z (D), 1W37 (E), 1O5K (F). The subfamily specific residues Ala135 in protein 1W3 (KDGA) and Leu141 in 1F5Z (NAL) shows a larger percentage of BSA compared to ASA compared to other residues in that segment. In DHDPS 1O5K, the catalytic signature residue Arg137 show a larger percentage of BSA compared to ASA. The next residue Thr138 is totally buried. Similar phenomenon is seen in protein 3B4U, that lacks signature arginine but possesses V144 whose BSA is larger than ASA and the next residue Thr145 is totally buried. Residues in protein 3E96 are largely accessible, with Glu148 showing a relatively larger BSA compared to other residues

### Major structural motions

The structural stability of the protein of respective family was assessed by Random mean square deviation (RMSD), Random mean square fluctuations (RMSF) and radius of gyration R_g_ which is an indicator for protein compactness (The details are discussed in Supplementary information SI6: SF5-SF6). The PCA results that describe the major motion in all the four proteins showed many motion vectors in the protein 3E96 indicate that it is highly dynamic, compared to the other three proteins (Vectors showing major motions, more particularly the interfacing segments, are shown in supplementary information SI6: SF7-SF10).

In 3E96, the vectors indicating major motions demonstrated that loop B and loop B1 drifts from loop A and moves towards the periphery. The plot showing a change in distance with respect to 50ns time of simulation indicates that there is a substantial increase in distance as the segments loop B, and loop B1 drifts from loop A compared to crystal structure (Figure 8A& B; Black spectra). The drifting of loop B1 and loop B from loop A opened the interface subset IS1. The motion vectors also demonstrated that loop A2 moves towards loop B and loop B1, which is evident from the decrease in distance between the segments compared to crystal structure (Figure 8C&D; Black spectra). The closer approach of loop A2 towards loop B1 and loop B strengthen the interface subset IS2. Vectors showing major motions, more particularly the interfacing segments and the structural analysis on the 0^th^, 49^th^ and 99^th^ frame, showing the wide opened IS1 and the tightened IS2 region is shown in the supplementary information (SI6: SF7-SF8)

**Figure 8:**
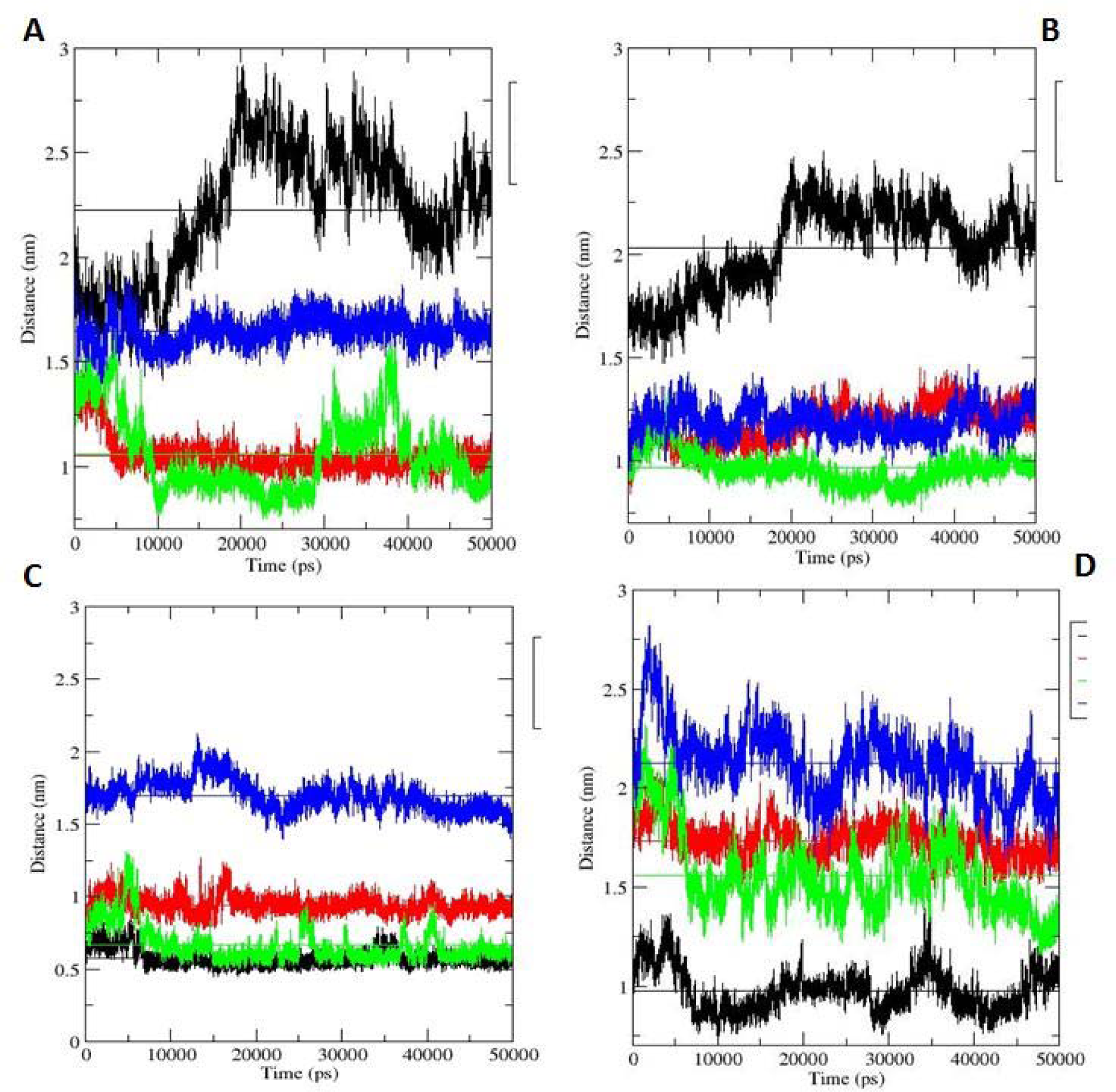
Change in distance between the segments with respect to time in the MD simulation. Curve colored in black (3E96), Green (1W3K), Red (2V8Z), Blue (1NAL). (A) Change in distance between Loop A/Helix A and Loop B/Helix B. The distance between the segments increased in protein 3E96, while it stabilized in rest of the proteins. (B) Change in distance between Loop A/Helix A and left end of loop B1. The distance between the segments increased in protein 3E96, while it stabilized in rest of the proteins. (C) Change in distance between Loop A2/helix A2 and Right end of loop B1. In Proteins 3E96 and 1W3K, the segments moves slightly closer compared to crystal structure, while the distance is stabilized throughout the simulation in protein YagE (2V8Z) and NAL (1NAL). (D) Change in distance between Loop A2/Helix A2 and Right end of Helix B/Helix B. Segments move closer in proteins 3E96 and KDGA (1W37), while the distance is stabilized in YagE (2V8Z). In NAL, the distance is more than the average during the major portion of simulation

In KDGA, PCA analysis showed that segments Helix A and B move towards the core that brought them closer initially and eventually drifts away and moves towards the periphery (Figure 8A; Green spectrum). Thus the two segments exhibit a sliding motion with respect to each other. The motion of helix A towards the core also brought it closer to static loop B1 compared to crystal structure which is maintained throughout the simulation (Figure8B; Green spectrum). The decreased distance between helix A and loop B1 strengthen the interface subset IS1. Few motion vectors seen at the right end of the relatively static loop B1 indicates that it moves in parallel to that of the motion of loop A2 and therefore there is almost no change in distance except for a slight decrease compared to crystal structure (Figure 8C; Green spectrum). However, the slight reduction in distance is highly insignificant leaving the IS2 unperturbed. Even though the motion of helix B towards the periphery brought it closer to helix A2, as evident from the decreased distance between the segments compared to crystal structure (Figure 8D; Green spectrum) the segments are not close enough to generate new interface region. Thus the interface segment IS2 is largely unaffected throughout the simulation period. Another significant observation is that the motion vectors at the extreme right end of helix B are pointed towards the core, indicating that this segment moves towards the core. Vectors showing major motions, more particularly in the interface region and the comparison of 0^th^, 49^th^ and 99^th^ frames illustrating the maintained tightness of IS1 region and the unperturbed IS2 segment is shown in Supplementary information (SI6: SF9-SF10)

PCA analysis on NAL protein showed that the right and the left end of loop A exhibits a twisting motion keeping the middle interfacing segment as the fulcrum. Vectors showing the major motions, more particularly in the interface segment are shown in Supplementary information (SI6: SF11). This twisting motion pushes the middle segment towards the core. The motion vectors at the right end of the loop B indicate its movement towards the core. Thus the middle interfacing segment of loop A and loop B are brought closer in simulation compared to crystal structure (Figure 8A; Blue spectrum). Also the static middle and left end of loop B1 maintains a constant distance from loop A (Figure 8B; Blue spectrum) indicating that IS1 region, as depicted through comparison of frames, is not opened but tightened compared to crystal structure (Supplementary information: SI6: SF12). Motion vectors indicate that the right end of loop B1 moves towards the core, while loop A2 moves antiparallel towards the periphery. Throughout the simulation, there is no significant change in the distance between the segments loop A2 and loop B1 (Figure 8C; Blue spectrum) indicating that the opened IS2 region in the crystal structure is maintained throughout the simulation period (Supplementary information: SI6: SF12). The movement of the right end of helix B towards the core as depicted by motion vectors is manifested as an increase in its distance from loop A2 (Figure 8D; blue spectrum). This observation indicated that the two segments do not come close to make surface contact as seen in protein 3E96. The increasing gulf between the surfaces of loop B1 and loop B also demonstrates that the latter moves towards the core in an ‘opening the lid’ fashion (Supplementary information: SI6: SF12)

In YagE, the only significant movement is the twisting motion of helix A with respect to helix B that holds each other from drifting away. Vectors showing the major motions, more particularly in the interface segment are shown in Supplementary information (SI6: SF13). Compared to the crystal structure, after 5ns, there is a decrease in distance between the segments which is maintained till 50ns simulation (Figure 8A; red spectrum). The decrease in distance tightens the IS1 region. Also, there is no major motion seen in loop B1. However, the slight increase seen in the distance between the segments loop B1 and Helix A is due to the twisting motion of the latter (figure 8B; red spectrum). Though this may slightly open up the IS1 region, comparison of frames showed that the closing of helix A and helix B maintain the steric crowding of IS1 (Supplementary information: SI6: SF14). The distance between helix A2 and the right end of loop B1 is maintained as seen in crystal structure throughout the simulation period as there is no major motion in the constituent segments (Figure 8C; red spectrum). Also, the distance between the right end of helix B and helix A2 in the crystal structure is maintained throughout the simulation period (Figure 8D; red spectrum). Thus there is no perturbation of the IS2 throughout the simulation period (Supplementary information: SI6: SF14)

Time-dependent analysis of the cavity volume of the four proteins measured by EPOCK**^73^** substantiated our PCA results. From the start of the simulation, the cavity volume in 3E96 keeps increasing because of the observed drifting motion of loop A and B with respect to each other towards the periphery. In other three proteins, the cavity volume is mostly unaffected due to the sliding and twisting motion of the IS1 forming segments in KDGA, NAL and YagE, that keeps the cavity intact. (Method discussion and figure depicting the change in the volume with simulation time is given in supplementary information SI6: SF15)

## Discussion

With pyruvate being the primary substrate of NAL superfamily proteins, logically NAL superfamily proteins should have evolved for binding pyruvate first and then should have diverged to perform subfamily specific enzymatic functions. Proteins 3E96 and 3D0C though retain the scaffold of NAL superfamily, lack the essential catalytic residues to bind pyruvate and therefore represent an inactive and possibly the earliest form of the superfamily. Next stage of evolution developed proteins that can bind pyruvate but cannot perform subfamily specific functions, Proteins 3EB2 and 3B4U retain the scaffold as well as the residues essential for binding pyruvate but lack the subfamily-specific residues harbored on the segment trailing the beta strand 5, thus represents an intermediate form. Later stage of evolution developed proteins that perform different functions of NAL, DHDPS, and KDGA.

Active site cavity topography analysis differentiated inactive form and active forms viz., a large cavity with the elongated topology in the former and trimmed cavity with the side-wise tilted topology in the latter. The transition from elongated to side-wise tilted topology is distinctly sharp as the tilted topology is demonstrated for intermediate forms as well. While the large elongated cavity in inactive forms spread across both the monomers of the biological dimer, the side-wise tilted topology in the active forms including the intermediates is restricted within the monomer resulting in two active site cavity, one for each monomer. This observation illustrates an evolution design of ‘randomness to order’ and ‘chopping of not-so-useful structural segments’

The strength of Interfacial regions IS1 and IS2 is the primary determinant of the active site cavity topology. Analysis of interfacial regions IS1 and IS2 contouring the active site cavity established tight IS2 and loose IS1 in inactive forms compared to opposing scenario in the active forms. In the inactive forms, the IS1 is weak and insignificant. The mild surface contact between the side chains of Lys145 harbored on loop A, and Pro118 on loop B1 in depth do not close the cavity but remain a backbone component of the cavity. Moreover, the segment loop B is at a more significant distance from loop A and oriented towards the outer region of the protein. These structural features provided the needed space for the cavity to spread further across the monomer B, thus resulting in a single cavity with elongated topology. The considerable distance between loop A and loop B is the reason for the broader head of the cavity. The tighter IS2 region restricts the cavity which is manifested as the narrow tail. In rest of the proteins, the segments Loop A/helix A and left end of Loop B1 are so near to each other and have their surfaces buried giving rise to strong and tighter IS1. Also, the Loop B/helix B is oriented towards the core and are positioned so close to Loop A/helix A, adding more steric hindrance around already tighter IS1. This structural feature restricts the cavity from spreading across the monomer B. Thus distinct cavities should have arisen for each monomer of biological dimers.

Orientation and participation of loop B/helix B provides a strong hint for the evolution of active forms. The already tighter IS2 region formed by loop B1 and loop A2 is further tightened by the partaking of Loop B in the inactive forms. The orientation of loop B especially the trough topology segment of Pro147 and Glu148, towards the outer regions of the protein facilitate this segment to partake in the formation of IS2 along with loop B1 and loop A2. The participation of Glu148 and Asp146 in the formation of IS2 is evident from the relatively larger buried surface area of these residues compared to other residues in the segment. In the active forms as the loop B/helix B is shifted towards the core and was positioned close to loop A/helix A and moreover the segment in loop B/helix B that is in alignment with Pro147 and Glu148 of 3E96 exhibiting crest topology consequently results in large distance from loop A2/helix A2. Thus loop B/helix B do not partake in the formation of IS2. However, they take part in IS1 or add steric hindrance around already tighter IS1 because of its proximity to loop A/helix A. Shifting of loopB/helix B towards the core makes the IS2 relatively open and this opened interfacial region should have evolved as the possible entry channel for the substrates to enter the active site cavity. The shallow concave surface morphology of IS2 formed by the right end of loop B1 and loop A2/helix A2 is unrestricted which may enable the entry of the substrates in the active proteins. Also, the positive coulombic potential of IS2 region would attract negatively charged pyruvate. In the inactive forms, though the buried surfaces of loop B1 and loop A2 forms a shallow concave surface, the buried surfaces of loop A2 and trough segment of loop B make the IS2 region sterically hindered that may restrict the substrates to enter the protein. In addition to steric hindrance, the negative coulombic electrostatic potential of IS2 region would repel electronegative charged pyruvate. We argue that the evolution framework should have made the inaccessible IS2 seen in the inactive forms to accessible IS2 in the active forms by executing the shift of loop B from outer regions towards the core. The shifted loop B opened the IS2 but in turn made the IS1 region inaccessible, which affected the change in cavity topography from elongated topography in the inactive forms to side-wise tilted topography in the active forms.

PCA analysis demonstrated that in protein 3E96, the loose IS1 and tight IS2 seen in the crystal structure is maintained or even enhanced in the simulation. This indicates there is no opening of the IS2 region throughout the simulation period that might make it active. In rest of the proteins, the already tighter IS1 and relatively loose IS2 seen in the crystal structure is maintained or enhanced in the simulation indicating that there is no open up of IS1 region or closure of entry channel IS2 region that may mediate a shift in cavity topography from sidewise tilted to elongated one to make the proteins inactive. Summarily, the opposing interface scenario in active and inactive forms and their conformational stability vis-à-vis., the unchanged scenario in their interfacial regions IS1 and IS2, indicates the handwork of evolution in the emergence of stable divergent active forms from an inactive form which itself is stable. Motion vectors and the movement of loop B in the ‘opening the lid’ fashion towards the core in protein 1NAL and motion vectors at the extreme right end of protein 1W37 (KDGA) substantiates the claim that loop B moved from the outer regions towards the core as active forms evolved.

Opposing trend vis-à-vis the number of occupants of loop A/helix A (loop B/helix B) and loop A2/helix A2 in the inactive and the active forms suggests evolutionary implications. Large numbers of residues in helix A (helix B) in the active forms should have been a result of later evolved insertions which consequently resulted in a helical secondary architecture compared to loop architecture in the inactive forms which harbor a lesser number of residues. The inserted residues contributed to the tightening of IS1, which is evident from the larger percentage buried surface area of this segment when compared to the percentage BSA of loop A in the inactive forms. The later evolved insertions in loop A/helix A included the subfamily specific residue viz., arginine of DHDPS, alanine of KDGA and leucine of NAL and the strictly conserved threonine/serine next to it. SASA analysis showed that the signature residues and their highly conserved immediate neighbor threonine/serine are primarily buried and are significant contributors to IS1. Thus these residues not only strengthen IS1 but also decide the subfamilial classifications indicating that these residues should have evolved much later. The evolved insertion process also resulted in the replacement of Asp146 and Glu148, the residues that contribute to the electronegativity to IS2 in protein 3E96. The fewer residues in the loop A(loop B) of the inactive forms not only lack the signature residues but also the threonine/serine marker thereby clearly showing them out of the league of the active forms.

With respect to segment loop A2/helix A2, the inactive forms harbor a more significant number of residues that contribute to IS2. In the active forms, the deletions in the segment which could have possibly evolved later results in weakened IS2. Deletion in loop A2/helix A2 and insertion in loopA/helix A should be excellent handwork of evolution, as the former is necessary to open the entry channel, while the latter is necessary for tightening the IS1 that consequently should have tilted the cavity topology towards the entry channel. The insertion and deletions also seemed to have a role to play in the orientation of loop B/helix B. Shortened loop A2/helix A2 due to deletions consequently limited it to form surface contact with loop B/helix B that eventually restricted loop B/helix B to be a part of IS2. The ‘freed’ loop B/helix B segment as itself has evolved to add insertions moved towards the core to take part or to provide steric crowding around IS1. Thus destabilization of IS2 due to deletions in loop A2/helix A2 is adequately compensated by stabilization of IS1 due to insertion in loop A/helix A and loop B/helix B, that consequently resulted in the evolution of active enzymes.

## Conclusion

As we attempted to correlate the variation in the cavity topology to the divergence in the activity among the subfamilies, we discovered that the modern day NAL superfamily proteins should have evolved from inactive ancestral forms. On an ancestral scaffold, random mutagenesis led to the cavity forming deletions at the secondary segment trailing the alpha helix 10 which is adequately compensated by cavity filling insertions in segment trailing beta strand 5. These changes which we propose as an evolutionary framework weakened the interface subset IS2, that triggers the movement of the secondary segment trailing the beta strand 5 towards the core, and complemented by cavity filling insertions led to the formation of tight interface subset (IS1). Tight IS1 and loose IS2 seen in evolved proteins as opposed to loose IS1 and tight IS2 seen in the inactive forms may have designed the entry channel and affected the change in the cavity topology. The evolved side-wise tilted topology is catalytically relevant as it is inclined towards the entry channel, thus justifying the emergence of active modern-day NAL proteins.

