## Supplementary Table ST1 for "Evolutionary significance of structural segments trailing beta strand 5 and alpha helix 10 in the design of catalytically active NAL superfamily"

| PDBid | Subfamily |
| --- | --- |
| 3E96 | DHDPS |
| 3EB2 | DHDPS |
| 2R8W | DHDPS |
| 3D0C | DHDPS |
| 3DI0 | DHDPS |
| 3B4U | DHDPS |
| 1W37 | KDPGA |
| 2NUW | KDPGA |
| 1F5Z | NAL |
| 1NAL | NAL |
| 2V8Z | YagE (KDGA) |
| 1DHP | DHDPS |
| 1O5K | DHDPS |
| 3A5F | DHDPS |
| 3G0S | DHDPS |
| 1XXX | DHDPS |
| 3H5D | DHDPS |
| 3NOE | DHDPS |
| 1XL9 | DHDPS |
| 3CPR | DHDPS |
| 3M5V | DHDPS |
| 2RFG | DHDPS |
| 3FLU | DHDPS |
| 2EHH | DHDPS |

**Supplementary information SI1**

**Supplementary table ST1:** The PDB Ids and the protein name are given

**Supplementary figure SF1: Naming and Definition**

**Figure SF1:** **(A)** Secondary structural segments that are involved in formation of interfacial region in the biological dimer of the protein 3E96. Image **(B)** represents the corresponding segments in rest of the proteins. In some proteins such as 1NAL, it is loop A instead of helix A and helix B. Monomer A is coloured yellow, while monomer B is coloured red. Among the segments, loop/Helix A (Blue coloured), loop B/helix B (Blue coloured), loop B1 (Green) and Loop A2/helix A2 (Cyan coloured) are considered for analysis.

**Supplementary information SI2 (Interface analysis)**

**Percentage Buried Surface area**

Buried surface area of the interfacial region contouring the active site cavity was done using PDBePISA. The methodology is discussed in the materials and methods section. The percentage buried surface area of IS1 region with respect to the total is less than the IS2 region in proteins 3E96 and 3D0C, while in rest of the proteins, percentage BSA of IS1 is larger than IS2 region. This indicates that for proteins 3E96 and 3D0C, the IS2 has a stronger hydrophobic effect compared to IS1, while it is the opposite in rest of the proteins.

|  | BSA_IS1% | BSA_IS2% |
| --- | --- | --- |
| 3 E96 | 10.1 | 89.2 |
| 3D0C | 7.3 | 92.4 |
| 1F5Z | 85.7 | 14.3 |
| 1O5K | 74.6 | 25.4 |
| 1NAL | 77 | 23 |
| 1W37 | 52.3 | 47.7 |
| 1XL9 | 80 | 20 |
| 1XXX | 96 | 4 |
| 2EHH | 75 | 25 |
| 2NOE | 74 | 26 |
| 2NUW | 57 | 43 |
| 2R8W | 74.2 | 25.8 |
| 2R91 | 48.5 | 51.5 |
| 2RFG | 81.2 | 18.8 |
| 2V8Z | 49.3 | 50.7 |
| 3AF5 | 63.7 | 36.3 |
| 3B4U | 88.3 | 11.7 |
| 3CPR | 72.4 | 27.6 |
| 3DI0 | 70 | 30 |
| 3EB2 | 71.6 | 28.4 |
| 3FLU | 78 | 22 |
| 3G0S | 72.8 | 27.1 |
| 3H5D | 72.37 | 27.63 |
| 3M5V | 71.2 | 28.8 |

**Figure SF2:** Chart showing protein 3E96 and 3D0C have a larger percentage of buried surface area for IS2 (Brown bar) compared to IS1 (Blue bar), while the rest of the proteins have a larger area for 1S1 compared to IS2.

**Supplementary ST2:** Table listing the numerical values of percentage BSA of IS1 and IS2, deduced from total BSA (Å^2^) and ASA (Å^2^).

| **PDBid** | **Total ASA**(Å^2^) | **Total BSA**(Å^2^) | **Total BSA% wrt ASA** | **BSA_IS1**(Å^2^) | **BSA_IS1%** | **BSA_IS2**(Å^2^) | **BSA_IS2%** |
| --- | --- | --- | --- | --- | --- | --- | --- |
| 3E96 | 2115.7 | 239.15 | 11.3 | 24.25 | 10.1 | 214.9 | 89.2 |
| 3D0C | 2408.8 | 252 | 10.4 | 18.6 | 7.3 | 233.4 | 92.4 |
| 1F5Z | 2356.7 | 188.85 | 8.0 | 161.85 | 85.7 | 27 | 14.3 |
| 1O5K | 1995.45 | 304.3 | 15.24 | 226.85 | 74.6 | 77.45 | 25.4 |
| 1NAL | 2298.75 | 148.4 | 6.4 | 114.4 | 77 | 34 | 23 |
| 1W37 | 2277.4 | 303.65 | 13.3 | 158.75 | 52.3 | 145 | 47.7 |
| 1XL9 | 1976.95 | 317.55 | 16 | 254 | 80 | 64 | 20 |
| 1XXX | 1878.7 | 228 | 12.13 | 218.35 | 96 | 9.65 | 4 |
| 2EHH | 1804.55 | 341.5 | 19 | 255.75 | 75 | 85.75 | 25 |
| 2NOE | 1944.3 | 344.2 | 17.7 | 254.3 | 74 | 90 | 26 |
| 2NUW | 2267.7 | 287.55 | 12.7 | 163.8 | 57 | 123.75 | 43 |
| 2R8W | 2204.1 | 361.25 | 16.4 | 268 | 74.2 | 93.25 | 25.8 |
| 2R91 | 2266.55 | 292.95 | 13 | 142.25 | 48.5 | 150.7 | 51.5 |
| 2RFG | 2059.35 | 334.1 | 16.2 | 271.2 | 81.2 | 62.9 | 18.8 |
| 2V8Z | 2204.75 | 355.6 | 16.1 | 175.25 | 49.3 | 180.35 | 50.7 |
| 3AF5 | 1843.45 | 314.9 | 17 | 200.45 | 63.7 | 114.45 | 36.3 |
| 3B4U | 2048.9 | 202.5 | 9.9 | 178.85 | 88.3 | 23.65 | 11.7 |
| 3CPR | 3875.5 | 210.25 | 5.4 | 152.3 | 72.4 | 57.95 | 27.6 |
| 3DI0 | 2037.45 | 368.2 | 18 | 258.1 | 70 | 110.1 | 30 |
| 3EB2 | 2446.15 | 305.4 | 12.5 | 218.55 | 71.6 | 86.85 | 28.4 |
| 3FLU | 1955.5 | 315.8 | 16.14 | 246.35 | 78 | 69.45 | 22 |
| 3G0S | 2023.5 | 317.15 | 15.7 | 230.95 | 72.8 | 86.2 | 27.1 |
| 3H5D | 2084 | 328 | 15.73 | 237.4 | 72.37 | 90.6 | 27.63 |
| 3M5V | 1976.2 | 318.3 | 16.1 | 226.6 | 71.2 | 91.7 | 28.8 |

**Supplementary information SI3**

**Solvation Free energy Gain due to Interface formation (∆^i^G, kcal/mol)**

**Δ^i^G** indicates the solvation free energy gain upon formation of the interface, in kcal/mole. The value is calculated as difference in total solvation energies of isolated and interfacing structures. Negative **Δ^i^G** corresponds to hydrophobic interfaces, or positive protein affinity. Protein 3E96 and 3D0C shows negative free energy gain due to formation of IS2 compared to positive gain for the formation of IS1, indicating stronger interactions in IS2 and weaker ones in IS1. In rest of the proteins, except for few exceptions, formation of IS1 show a larger negative free energy gain compared to IS2, indicating stronger interactions in IS1 and weaker ones in IS2. The values do not include hydrogen bonding and salt bridges. The exceptions are justifiable as, though the interactions for IS2 is stronger, the shallow concave morphology, needed for unhindered transport of substrate into active site, is unperturbed and therefore it is not a cause for concern.

|  | deltaG_IS1 | deltaG_IS2 |
| --- | --- | --- |
| 3D0C | 0.5 | -2.9 |
| 3 E96 | 0.6 | -3.2 |
| 1F5Z | -2.7 | -0.3 |
| 1NAL | -1.7 | -1.1 |
| 1W37 | -2.6 | -0.9 |
| 1XL9 | -2.3 | -0.7 |
| 1XXX | -1.2 | -0.1 |
| 2EHH | -5.2 | -2 |
| 2NOE | -0.3 | -0.3 |
| 2NUW | -2 | -2.4 |
| 2R8W | 0.3 | -0.4 |
| 2R91 | -1.8 | -1.5 |
| 2RFG | -0.8 | -0.7 |
| 2V8Z | -2.5 | -3.8 |
| 3A5F | -0.2 | 2.9 |
| 3B4U | -4 | -0.5 |
| 3CPR | -0.8 | 0.2 |
| 3DI0 | 0.9 | -2.1 |
| 3EB2 | 1.3 | -1.6 |
| 3FLU | -2.3 | -0.7 |
| 3G0S | -1.1 | -1.2 |
| 3H5D | -2.4 | -0.5 |
| 3M5V | -1.3 | -1.8 |
| 3NOE | -0.3 | -0.3 |
| 105K | -1 | -1.7 |

**Figure SF3:** Chart showing protein 3E96 and 3D0C show a negative free energy gain ((∆^i^G, kcal/mol) due to interface formation of IS2 compared to IS1, indicating stronger IS2 and weaker IS1. Rest of the proteins, but for few exceptions, show a larger negative gain for the formation of IS1 compared to IS2, indicating stronger IS1 and weaker IS2.

**Supplementary table ST3:** Table listing the numerical values of solvation free energy gain (∆iG, kcal/mol) due to interface formation of IS1 and IS2

| **PDBid** | **Solvation free energy gain due to interface formation of IS1**  **∆G, kcal/mol** | **Solvation free energy gain due to interface formation of IS2**  **∆G, kcal/mol** |
| --- | --- | --- |
| 3E96 | 0.6 | -3.2 |
| 3D0C | 0.5 | -2.9 |
| 1F5Z | -2.7 | -0.3 |
| 1O5K | -1 | -1.7 |
| 1NAL | -1.7 | -1.1 |
| 1W37 | -2.6 | -0.9 |
| 1XL9 | -2.3 | -0.7 |
| 1XXX | -1.2 | -0.1 |
| 2EHH | -5.2 | -2 |
| 2NOE | -0.3 | -0.3 |
| 2NUW | -2 | -2.4 |
| 2R8W | 0.3 | -0.4 |
| 2R91 | -1.8 | -1.5 |
| 2RFG | -0.8 | -0.7 |
| 2V8Z | -2.5 | -3.8 |
| 3AF5 | -0.2 | 2.9 |
| 3B4U | -4 | -0.5 |
| 3CPR | -0.8 | 0.2 |
| 3DI0 | 0.9 | -2.1 |
| 3EB2 | 1.3 | -1.6 |
| 3FLU | -2.3 | -0.7 |
| 3G0S | -1.1 | -1.2 |
| 3H5D | -2.4 | -0.5 |
| 3M5V | -1.3 | -1.8 |

**Supplementary information SI4: Active site cavity analysis**

**Active site cavity mapping and volume calculation**

Active site volume of all the proteins mentioned in ST1 was measured using CastP (<http://sts-fw.bioengr.uic.edu/castp/calculation.php>) (Dundas et al., 2006). NAL superfamily proteins are active as dimers,. A tyrosine from the neighboring monomer (monomer partner of the biological dimer) takes part in the catalytic triad which also includes a strictly conserved tyrosine and a serine or threonine. Therefore, the calculations were made for dimers. Calculations were made for all the available biological dimers in each protein. CastP lists the entire cavity present in the proteins with details of volume, area and the residues present in the cavity. The cavity which lists all the catalytic residues- like the lysine that is involved in schiff base with pyruvate, the catalytic triad residues of serine/threonine, tyrosine and another tyrosine from the neighboring monomer- all the residues that deal with pyruvate and the secondary catalytic residues that are involved in hydrogen bonding with the other substrate, are taken as the appropriate active site cavity. Probe radius was set at 2.0Å, since, at this probe radius, the active cavity detected has accommodated in it almost all the catalytic residues in many of the proteins.

**Supplementary table ST4:** Table listing the active site cavity volume and surface area

| **Protein**  **PDBid** | **Subfamily** | **Species** | **Gram staining** | **Volume of the cavity (Å^3^)** | **Surface Area of the cavity (Å^2^)** |
| --- | --- | --- | --- | --- | --- |
| 3E96 | DHDPS (inactive) | *B. clausii* | Gram +ve | 1820.0/470.8 | 1004.0/441.8 |
| 3B4U | DHDPS  (inactive) | *A. tumifaciens* | Gram -ve | 1220.6/835.8 | 803.0/590.8 |
| 3EB2 | DHDPS (inactive) | *R. plaustris* | Gram -ve | 1195.1/860.0  658.3/729.2 | 719.1/450.2  382.7/391.5 |
| 1NAL | NAL | *E.coli* | Gram -ve | 998.6/1016.7  711.2/367.6 | 566.1/584.1  423.3/230.5 |
| 1F5Z | NAL | *H. influenza* | Gram -ve | 911/538.0  1007.4/964.3 | 524/326.6  547.3/526.7 |
| 1W37 | KD(P)GA | *S. sulfataricus* | Arachea | 840.3/693.1  666.9/684.7 | 566.8/534.8  484.4/492.5 |
| 1XXX | DHDPS | *M. tuberculosis* | Gram +ve | 749.7/735.5  758.6/761.2  787.5/739.5  767.7/768.8 | 438.7/439.4  449.4/442.7  458.5/441.9  448.7/447.5 |
| 3D0C | DHDPS (inactive) | *O. iheyensis* | Gram +ve | 777.3/772.9 | 516.9/551.3 |
| 2NUW | KD(P)GA | *S. acidocaldaricus* | Arachea | 697.6/635.1 | 560.5/548.5 |
| 3DI0 | DHDPS | *S. aureus* | Gram +ve | 687.0/620.6 | 362.3/380.8 |
| 2R8W | DHDPS (inactive) | *A. tumifaciens* | Gram -ve | 610.5/573.7 | 385.5/434.9 |
| 1O5K | DHDPS | *T. maritima* | Gram -ve | 396.3/305.8  (Pyr removed) | 324.2/260.1 |
| 3NOE | DHDPS | *P. aerugenosa* | Gram -ve | 382.4/158.7 | 283.6/129.5 |
| 3H5D | DHDPS | *S. pneumonia* | Gram +ve | 335.3/285.4 | 263.0/234.6 |
| 3M5V | DHDPS | *C. jejuni* | Gram -ve | 316.4/269.2  311.4/275.1 | 238.8/212.3  259.6/215.4 |
| 2RFG | DHDPS | *H. chejuensis* | - | 290.9/284.9  285.3/287.3 | 239.5/230.4  232.9/231.2 |
| 1XL9 | DHDPS | *B. anthracis* | Gram +ve | 244.1/232.8  275.9/235.9 | 187.4/213.2  207.0/189.4 |
| 2EHH | DHDPS | *A.aeolicus* | Gram -ve | 252.8/127.3  190.6/111.3 | 254.0/164.3  212.7/136.5 |
| 3A5F | DHDPS | *C. botulinum* | Gram +ve | 241.5/240.7 | 216.0/214.6 |
| 3G0S | DHDPS | *S. typhimurium* | Gram -ve | 230.0/208.9 | 184.7/172.6 |
| 2V8Z | YagE | *E.coli K12* | Gram -ve | 201.1/195.9  205.1/203.5 | 181.3/177.9  192.1/188.6 |
| 3CPR | DHDPS | *C. glutamicum* | Gram +ve | 134.3/133.5 | 131.2/130.4 |
| 3FLU | DHDPS | *N. meningtidis* | Gram –ve | 109.9/114.5  115.7/115.0 | 98.1/103.5  103.3/103.2 |

**Supplementary table ST5:** The constituent residues of the active site that were categorized as catalytic, non catalytic residues and residues from neighbouring monomer are listed.

| **PDBid** | **Catalytic residues** | **Non catalytic residues** | **Residues from neighboring monomer** |
| --- | --- | --- | --- |
| 3E96 | N55, T56, Y143, G194, T195, A196, H117 | I18, M111, M114, P115, I116, H117, P118, V120, D146, P147, E148, A169, I170, N171, K198, W199, T212, S213, G214, E249, N256, Q257, N260 | M114, P115, I116, H117, P118, Y119, V120, K145, D146, P147, E148, N256 |
| 3B4U | T46, T47, Y139, V144, K169, G193, D194, E195 | A10, S143, S171, S172, G173, R196, I210, S211, G212, V213, V235, V238, V239, L241, L242, P245, V246, T247, P248, V250, L270, V271 | F110, N112, V113, S114 |
| 3EB2 | S47, Y136, A189, S190, A191, Y110, F141 | Y11, N138, Q140, A166, T168, W240, N243, E244, F246, A247, K248, F249, N250, L251, Q273, A274, L276, T277, E280 | F111, P112, L113, Q117 |
| 1NAL | S47, T48, Y137, K165, G189, Y190, D191, E192, S208 | A11, L142, T167, G207, T209, I243, K248, T249, G250, V251, F252, R253, F274, G275, P276, D278, K280, Y281 | P112, S114, E116, E117 |
| 1W37 | T43, T44, Y130, K155, T157, G179, A198, S241, N245 | P7, T134, I158, E159, S180, D181, V196, I233, S236, L242 | P105 |
| 1XXX | T54, T55, Y143, R148, G194, D195 | G147, A173, A197, V213, C248, M251, R253, L254, G255, G256 | P120, P121, Q122, V151, P152, E154 |
| 3D0C | N53, T54, Y141, K165, G192, T193, A194 | I16, V49, G52, V83, G85, I110, H111, I139, N169, K196, W197, T210, S211, G212, E247, A251, N255, N258 | NIL^*^ |
| 3DI0 | T46, T47, Y135, R140, K163, G189, N190, D191 | S139, A165, D192, V208, L241, L244, D247, I248, N249 | NIL^*^ |
| 2R8W | S77, T78, Y166, T171, K194, S223, G224 | F41, T170, T171, P196, L197, W226, Y240, V242, G284, S285, I286, R287 | T141, P142 |
| 3NOE | T44, T45, Y133, R138, K161, G186, E187, | G137, A163, T164, V205, S247, N248 | NIL^*^ |
| 3H5D | T50, T51, Y140, R145, K168, G192, E193 | G144, C170, V211, P253, S254 | NIL^*^ |
| 3G0S | T44, T45, Y133, R138, K161, G186, D187 | S137, A163, V205, N248 | NIL^*^ |
| 2V8Z | S56, G57, K174, T176, G202, Y203, A221, | P20, S220, Y259, F265, I269 | W120 |
| 3CPR | T59, T60, Y148, R153, G199, D200 | V218, Q256 | NIL |
| 3FLU | T44, T45, Y133, R138, K161, G186 | A163, V205 | NIL |

The appropriate cavity of the protein 3E96 constitutes residues including those that are harbored on the loop A1 from its own monomer and loop B2 from the neighboring monomer thus explaining large number of catalytically insignificant residues as its constituents. In rest of the proteins the residues harboured on the loop A1 and loop B2/helix B of neighboring monomer are not accommodated in their active site cavity. The cavity in 3E96 and rest of the proteins spread to include the residues from loop B1 of the neighboring monomer. However there is a marked difference in the constituent residues from this loop, which explains the uniqueness of the cavity of protein 3E96.

**Supplementary Information SI5: Accessible surface area analysis on the segment trailing beta strand 5**

Percentage ASA and BSA of the segment trailing beta strand 5 (loop A/helix A) calculated using AreaiMOL of CCP4 suite showed that proteins 3E96 and 3D0C have more than 80% of their surface area exposed to solvent, with former having 86% of their surface solvent exposed. All other proteins have their percentage ASA decreased when compared to 3E96 and 3D0C. However there is an exception in NAL subfamily, wherein the protein 1NAL show 85% of their surface solvent exposed. 1F5Z protein, which also belongs to NAL subfamily, has 80% of their surface solvent exposed. KDGA family proteins show further decrease in their percentage ASA. The proteins belonging to this family have 70 to 80% of their surface exposed to solvent. DHDPS proteins showed least percentage ASA compared to other subfamilies. They show very low percentage values that are distributed between 55 to 70%. This indicates that the segment in DHDPS proteins is largely buried followed by KDGA and NAL subfamilies. The segments in 3E96 and 3D0C are less buried and are largely solvent exposed.

|  | ASA% | BSA% |
| --- | --- | --- |
| 3 E96 | 86 | 14 |
| 3B4U | 70 | 30 |
| 1F5Z | 80 | 20 |
| 1W37 | 78 | 22 |
| 1O5K | 64 | 36 |
| 2V8Z | 72 | 28 |
| 3D0C | 82 | 18 |
| 1NAL | 85 | 15 |
| 3EB2 | 65 | 35 |
| 2R8W | 66 | 34 |
| 3CPR | 69 | 31 |
| 2NUW | 74 | 26 |
| 2R91 | 79 | 21 |
| 1XL9 | 59 | 41 |
| 3H5D | 60 | 40 |
| 1XXX | 64 | 36 |
| 3M5V | 62 | 38 |
| 2EHH | 61 | 39 |
| 3A5F | 63 | 37 |
| 3DI0 | 61 | 39 |
| 3G0S | 57 | 43 |
| 3NOE | 56 | 44 |
| 3FLU | 57 | 43 |
| 2RFG | 58 | 42 |

**Figure SF4:** Chart showing percentage ASA and BSA of the segment trailing beta strand 5 (loop A/helix A) in NAL superfamily proteins calculated using AreaiMOL of CCP4 suite.

**Supplementary table ST6:** The percentage ASA of monomer with respect to dimer and BSA with respect to the dimer of the segment trailing beta strand 5 in NAL superfamily proteins calculated using AreaiMOL of CCP4 suite are listed.

| **PDBid** | **ASADimer**(Å^2^) | **ASAMonomer**(Å^2^) | **ASA%** | **BSA**(Å^2^) | **BSA%** |
| --- | --- | --- | --- | --- | --- |
| 3E96 | 388.6 | 451.6 | 86 | 63 | 14 |
| 3D0C | 319.6 | 389 | 82 | 69.4 | 18 |
| 1NAL | 437.85 | 514.3 | 85 | 77 | 15 |
| 1F5Z | 485.7 | 601 | 80 | 115.3 | 20 |
| 1W37 | 370.6 | 478.5 | 78 | 108 | 22 |
| 1O5K | 271.1 | 425.7 | 64 | 154 | 36 |
| 2V8Z | 326.8 | 455.2 | 72 | 128.4 | 28 |
| 3EB2 | 422.6 | 647.05 | 65 | 224.45 | 35 |
| 2R8W | 402.5 | 611.8 | 66 | 209 | 34 |
| 3CPR | 331.2 | 478.7 | 69 | 147.5 | 31 |
| 2NUW | 334.2 | 450.6 | 74 | 116.4 | 26 |
| 2R91 | 388.6 | 492.3 | 79 | 103.7 | 21 |
| 1XL9 | 308.4 | 521.1 | 59 | 212.7 | 41 |
| 3H5D | 289.4 | 482.1 | 60 | 192.7 | 40 |
| 1XXX | 298.7 | 464.17 | 64 | 165.45 | 36 |
| 3M5V | 243.5 | 391.8 | 62 | 148.3 | 38 |
| 2EHH | 291.5 | 480.1 | 61 | 188.6 | 39 |
| 3A5F | 253.4 | 400.8 | 63 | 147.5 | 37 |
| 3DI0 | 312.3 | 513 | 61 | 200.7 | 39 |
| 3G0S | 248 | 434.5 | 57 | 186.5 | 43 |
| 3NOE | 260.35 | 460.9 | 56 | 200.55 | 44 |
| 3FLU | 255.2 | 446.1 | 57 | 190.9 | 43 |
| 2RFG | 295.25 | 511.15 | 58 | 215.9 | 42 |

**Supplementary table ST7:** The percentage ASA of monomer with respect to total area and BSA with respect to the total area for individual residues harboured on segment trailing beta strand 5 (loop A/helixA) in protein 3E96 are listed

| **Residues** | **Total Area**(Å^2^) | **ASA**(Å^2^) | **ASA%** | **BSA**(Å^2^) | **BSA%** |
| --- | --- | --- | --- | --- | --- |
| K145 | 106.7 | 85.9 | 81 | 20.8 | 19 |
| D146 | 41.7 | 37.4 | 90 | 4.3 | 10 |
| P147 | 81.3 | 81.3 | 100 | 0 | 0 |
| G148 | 160.6 | 122.7 | 76 | 37.9 | 24 |
| I149 | 3.8 | 3.8 | 100 | 0 | 0 |
| S150 | 57.5 | 57.5 | 100 | 0 | 0 |

**Supplementary table ST8:** The percentage ASA of monomer with respect to total area and BSA with respect to the total area for individual residues harboured on segment trailing beta strand 5 (loop A/helixA) in protein 3B4U are listed

| **Residues** | **Total Area**(Å^2^) | **ASA**(Å^2^) | **ASA%** | **BSA**(Å^2^) | **BSA%** |
| --- | --- | --- | --- | --- | --- |
| I141 | 15.8 | 14.2 | 90 | 1.6 | 10 |
| P142 | 27.9 | 27.9 | 100 | 0 | 0 |
| S143 | 90 | 81.7 | 91 | 8.3 | 9 |
| V144 | 86.8 | 20.8 | 24 | 66 | 76 |
| T145 | 11.6 | 0.1 | 0.8 | 11.5 | 99.1 |
| M146 | 144.8 | 88.1 | 61 | 56.7 | 39 |
| V147 | 4.1 | 4.1 | 100 | 0 | 0 |
| T148 | 72.3 | 72.3 | 100 | 0 | 0 |
| L149 | 1.4 | 1.4 | 100 | 0 | 0 |
| S150 | 20 | 20 | 100 | 0 | 0 |

**Supplementary table ST9:** The percentage ASA of monomer with respect to total area and BSA with respect to the total area for individual residues harboured on segment trailing beta strand 5 (loop A/helixA) in protein 1F5Z (NAL) are listed

| **Residues** | **Total Area**(Å^2^) | **ASA**(Å^2^) | **ASA%** | **BSA**(Å^2^) | **BSA%** |
| --- | --- | --- | --- | --- | --- |
| I138 | 17.9 | 16.1 | 90 | 1.8 | 10 |
| P139 | 25.5 | 25.5 | 100 | 0 | 0 |
| F140 | 181.9 | 147.4 | 81 | 34.5 | 19 |
| L141 | 109.6 | 34 | 31 | 75.6 | 69 |
| T142 | 20.5 | 17.1 | 83 | 3.4 | 17 |
| G143 | 48.6 | 48.6 | 100 | 0 | 0 |
| V144 | 24.9 | 24.9 | 100 | 0 | 0 |
| N145 | 123.2 | 123.2 | 100 | 0 | 0 |
| M146 | 17.2 | 17.2 | 100 | 0 | 0 |

**Supplementary table ST10:** The percentage ASA of monomer with respect to total area and BSA with respect to the total area for individual residues harboured on segment trailing beta strand 5 (loop A/helixA) in protein 1W37 (KDGA) are listed

| **Residues** | **Total Area**(Å^2^) | **ASA**(Å^2^) | **ASA%** | **BSA**(Å^2^) | **BSA%** |
| --- | --- | --- | --- | --- | --- |
| Y132 | 60.7 | 40.2 | 66 | 20.5 | 34 |
| P133 | 29.3 | 29.3 | 100 | 0 | 0 |
| T134 | 127.4 | 102.9 | 81 | 24.5 | 19 |
| A135 | 61.5 | 7.3 | 12 | 54.2 | 88 |
| T136 | 7.3 | 2.9 | 40 | 4.4 | 60 |
| G137 | 56.6 | 52.3 | 92 | 4.3 | 8 |
| K138 | 19.8 | 19.8 | 100 | 0 | 0 |
| D139 | 61.1 | 61.1 | 100 | 0 | 0 |
| I1.2 | 17.2 | 1.2 | 100 | 0 | 0 |

**Supplementary Table ST11:** The percentage ASA of monomer with respect to total area and BSA with respect to the total area for individual residues harboured on segment trailing beta strand 5 (loop A/helixA) in protein 1O5K (DHDPS) are listed

| **Residues** | **Total Area**(Å^2^) | **ASA**(Å^2^) | **ASA%** | **BSA**(Å^2^) | **BSA%** |
| --- | --- | --- | --- | --- | --- |
| V134 | 16 | 14.1 | 88 | 1.9 | 12 |
| P135 | 32.3 | 32.3 | 100 | 0 | 0 |
| G136 | 86.8 | 54.5 | 63 | 32.3 | 37 |
| R137 | 111 | 17 | 15 | 94 | 85 |
| T138 | 7.7 | 0 | 0 | 7.7 | 100 |
| G139 | 48.1 | 30 | 62 | 18.1 | 38 |
| V140 | 10.9 | 10.9 | 100 | 0 | 0 |
| N141 | 40.8 | 40.8 | 100 | 0 | 0 |
| V142 | 1.6 | 1.6 | 100 | 0 | 0 |

**Supplementary information SI6: Molecular Dynamics simulation**

**Structural stability**

The structural stability of the protein of respective family was assessed by Random mean square deviation (RMSD) and radius of gyration R_g_, a indicator for protein compactness (Figure SF5). The protein backbone RMSD calculated for 3E96 was initially found to raise from 1 Å to 6 Å, the trend continued up to 22 ns, after that the system stabilized. The backbone RMSD of YAGE, KDGA, and NAL were stable throughout the 50 ns simulation. Hence a 5-50 ns trajectories of 3E96 (inactive), YAGE, KDGA, and NAL used for further domain motion analysis using Essential dynamics. The structural compactness of the protein was ensured by the constant R_g_ value that is observed throughout the simulation.

The root mean square fluctuations (RMSF) calculated from MD simulations (Figure SF5) were converted to simulated B-factor (Figure SF6) using the equation given below.

$$B={\frac{8\pi}{3}}^{2}\times{(RMSF)}^{2}$$

The calculated and experimental B-factor was normalized by division of each B-factor value with the highest B-factor value found within a protein. A comparison between the two B-factors, reveal a high degree of correlation in its relative magnitude, this reiterate the MD simulation ability to reproduce the natural flexibility of the proteins in the investigation.

**Figure SF5:** Structural stability of 3E96 (inactive), YagE, KDGA and NAL. **A.** Root mean square deviation (RMSD) of back bone atoms relative to the starting structure during MD simulation. **B.** Radius of gyration (Rg) plot during the MD simulation. C. RMSF of the backbone atoms of individual residues.

|  | B-factor MD | B-factor X-ray |
| --- | --- | --- |
| 1 | 0.332693 | 0.746692 |
| 2 | 0.212622 | 0.635336 |
| 3 | 0.255504 | 0.575799 |
| 4 | 0.282514 | 0.562293 |
| 5 | 0.262545 | 0.503583 |
| 6 | 0.227343 | 0.460309 |
| 7 | 0.231055 | 0.455347 |
| 8 | 0.220174 | 0.43054 |
| 9 | 0.203661 | 0.393605 |
| 10 | 0.145033 | 0.389746 |
| 11 | 0.136841 | 0.340959 |
| 12 | 0.0989503 | 0.360529 |
| 13 | 0.120072 | 0.382029 |
| 14 | 0.12596 | 0.363561 |
| 15 | 0.146697 | 0.401874 |
| 16 | 0.163082 | 0.466373 |
| 17 | 0.191244 | 0.499449 |
| 18 | 0.225038 | 0.630099 |
| 19 | 0.281106 | 0.81505 |
| 20 | 0.305044 | 0.826902 |
| 21 | 0.293395 | 0.733186 |
| 22 | 0.22555 | 0.636439 |
| 23 | 0.232719 | 0.531422 |
| 24 | 0.195981 | 0.472161 |
| 25 | 0.195981 | 0.521499 |
| 26 | 0.205325 | 0.527288 |
| 27 | 0.22299 | 0.658765 |
| 28 | 0.18702 | 0.572216 |
| 29 | 0.165387 | 0.480981 |
| 30 | 0.193164 | 0.516262 |
| 31 | 0.191244 | 0.544101 |
| 32 | 0.165003 | 0.51957 |
| 33 | 0.165003 | 0.477122 |
| 34 | 0.191372 | 0.54796 |
| 35 | 0.184076 | 0.550441 |
| 36 | 0.178443 | 0.555954 |
| 37 | 0.200589 | 0.578556 |
| 38 | 0.220942 | 0.707277 |
| 39 | 0.223246 | 0.693219 |
| 40 | 0.252304 | 0.590408 |
| 41 | 0.176523 | 0.493936 |
| 42 | 0.186892 | 0.510474 |
| 43 | 0.153354 | 0.423374 |
| 44 | 0.135817 | 0.399394 |
| 45 | 0.100998 | 0.374311 |
| 46 | 0.12596 | 0.382856 |
| 47 | 0.12916 | 0.380099 |
| 48 | 0.159754 | 0.407387 |
| 49 | 0.179083 | 0.470232 |
| 50 | 0.159882 | 0.366593 |
| 51 | 0.15041 | 0.406836 |
| 52 | 0.144009 | 0.353363 |
| 53 | 0.172171 | 0.356946 |
| 54 | 0.19278 | 0.356395 |
| 55 | 0.203661 | 0.340684 |
| 56 | 0.200589 | 0.375689 |
| 57 | 0.240655 | 0.435502 |
| 58 | 0.244112 | 0.45011 |
| 59 | 0.253328 | 0.492558 |
| 60 | 0.21851 | 0.42613 |
| 61 | 0.191884 | 0.407663 |
| 62 | 0.216846 | 0.434399 |
| 63 | 0.225038 | 0.472161 |
| 64 | 0.186892 | 0.430265 |
| 65 | 0.188044 | 0.443495 |
| 66 | 0.221838 | 0.568633 |
| 67 | 0.21979 | 0.531698 |
| 68 | 0.182796 | 0.503308 |
| 69 | 0.196621 | 0.577453 |
| 70 | 0.235279 | 0.746968 |
| 71 | 0.22171 | 0.680265 |
| 72 | 0.201869 | 0.683572 |
| 73 | 0.228111 | 0.816428 |
| 74 | 0.24744 | 0.735667 |
| 75 | 0.228111 | 0.660695 |
| 76 | 0.182796 | 0.562018 |
| 77 | 0.173067 | 0.522326 |
| 78 | 0.156298 | 0.422547 |
| 79 | 0.131336 | 0.38065 |
| 80 | 0.13044 | 0.341235 |
| 81 | 0.115335 | 0.353363 |
| 82 | 0.12276 | 0.330485 |
| 83 | 0.129928 | 0.324421 |
| 84 | 0.146953 | 0.331312 |
| 85 | 0.179339 | 0.387541 |
| 86 | 0.186124 | 0.427784 |
| 87 | 0.21723 | 0.403528 |
| 88 | 0.204813 | 0.380375 |
| 89 | 0.171275 | 0.350606 |
| 90 | 0.187148 | 0.374587 |
| 91 | 0.211214 | 0.382304 |
| 92 | 0.193164 | 0.377619 |
| 93 | 0.176651 | 0.366593 |
| 94 | 0.210189 | 0.421444 |
| 95 | 0.220942 | 0.375413 |
| 96 | 0.192012 | 0.428886 |
| 97 | 0.207885 | 0.495314 |
| 98 | 0.24744 | 0.531974 |
| 99 | 0.241423 | 0.490077 |
| 100 | 0.228495 | 0.505788 |
| 101 | 0.185996 | 0.455623 |
| 102 | 0.178955 | 0.41731 |
| 103 | 0.145545 | 0.419239 |
| 104 | 0.12468 | 0.353638 |
| 105 | 0.0997184 | 0.344542 |
| 106 | 0.0999744 | 0.282249 |
| 107 | 0.104711 | 0.296582 |
| 108 | 0.162058 | 0.395259 |
| 109 | 0.206093 | 0.343991 |
| 110 | 0.258961 | 0.392503 |
| 111 | 0.240527 | 0.358324 |
| 112 | 0.257168 | 0.375413 |
| 113 | 0.180364 | 0.338203 |
| 114 | 0.217358 | 0.309537 |
| 115 | 0.233231 | 0.347023 |
| 116 | 0.240399 | 0.361907 |
| 117 | 0.249616 | 0.392503 |
| 118 | 0.211854 | 0.35667 |
| 119 | 0.177803 | 0.312293 |
| 120 | 0.18894 | 0.369901 |
| 121 | 0.202381 | 0.357222 |
| 122 | 0.169739 | 0.322216 |
| 123 | 0.140297 | 0.324146 |
| 124 | 0.167563 | 0.429162 |
| 125 | 0.194444 | 0.476571 |
| 126 | 0.162826 | 0.400221 |
| 127 | 0.151178 | 0.36301 |
| 128 | 0.188556 | 0.49118 |
| 129 | 0.214158 | 0.48677 |
| 130 | 0.197133 | 0.456725 |
| 131 | 0.186764 | 0.50634 |
| 132 | 0.174603 | 0.442944 |
| 133 | 0.14977 | 0.405182 |
| 134 | 0.1202 | 0.37817 |
| 135 | 0.103175 | 0.391951 |
| 136 | 0.0986943 | 0.411246 |
| 137 | 0.126344 | 0.356395 |
| 138 | 0.155786 | 0.327178 |
| 139 | 0.247312 | 0.364388 |
| 140 | 0.283922 | 0.304024 |
| 141 | 0.359191 | 0.402701 |
| 142 | 0.3702 | 0.439085 |
| 143 | 0.293523 | 0.353363 |
| 144 | 0.285842 | 0.38699 |
| 145 | 0.283538 | 0.409041 |
| 146 | 0.306068 | 0.431367 |
| 147 | 0.225422 | 0.361356 |
| 148 | 0.232335 | 0.296031 |
| 149 | 0.295955 | 0.40656 |
| 150 | 0.263697 | 0.440187 |
| 151 | 0.195725 | 0.392227 |
| 152 | 0.221454 | 0.434675 |
| 153 | 0.25704 | 0.521499 |
| 154 | 0.18702 | 0.51516 |
| 155 | 0.171147 | 0.485391 |
| 156 | 0.144265 | 0.459206 |
| 157 | 0.126728 | 0.372381 |
| 158 | 0.144009 | 0.415656 |
| 159 | 0.143881 | 0.421444 |
| 160 | 0.130312 | 0.341786 |
| 161 | 0.125832 | 0.331588 |
| 162 | 0.155146 | 0.314774 |
| 163 | 0.183436 | 0.318908 |
| 164 | 0.193292 | 0.298787 |
| 165 | 0.239247 | 0.295755 |
| 166 | 0.269713 | 0.32828 |
| 167 | 0.28149 | 0.350331 |
| 168 | 0.311956 | 0.320838 |
| 169 | 0.283922 | 0.349779 |
| 170 | 0.252944 | 0.328831 |
| 171 | 0.291731 | 0.348126 |
| 172 | 0.310164 | 0.388644 |
| 173 | 0.274066 | 0.381753 |
| 174 | 0.273169 | 0.416759 |
| 175 | 0.325781 | 0.490353 |
| 176 | 0.311828 | 0.539967 |
| 177 | 0.250128 | 0.685502 |
| 178 | 0.334485 | 0.864939 |
| 179 | 0.40809 | 0.957001 |
| 180 | 0.37276 | 0.940739 |
| 181 | 0.299283 | 0.918688 |
| 182 | 0.388249 | 0.838479 |
| 183 | 0.216974 | 0.628445 |
| 184 | 0.220814 | 0.460309 |
| 185 | 0.211726 | 0.388368 |
| 186 | 0.193036 | 0.36301 |
| 187 | 0.191628 | 0.33462 |
| 188 | 0.188684 | 0.369901 |
| 189 | 0.210829 | 0.341235 |
| 190 | 0.231823 | 0.369901 |
| 191 | 0.226446 | 0.392503 |
| 192 | 0.273938 | 0.358875 |
| 193 | 0.254864 | 0.34344 |
| 194 | 0.258065 | 0.373208 |
| 195 | 0.259217 | 0.351985 |
| 196 | 0.249744 | 0.343164 |
| 197 | 0.22299 | 0.338754 |
| 198 | 0.22299 | 0.399394 |
| 199 | 0.247952 | 0.450937 |
| 200 | 0.243216 | 0.462238 |

|  | B-factor MD | B-factor X-ray |
| --- | --- | --- |
| 1 | 0.926915 | 0.931912 |
| 2 | 0.78551 | 0.915406 |
| 3 | 0.721004 | 0.691197 |
| 4 | 0.71592 | 0.444292 |
| 5 | 0.509056 | 0.264787 |
| 6 | 0.304417 | 0.21458 |
| 7 | 0.268827 | 0.295048 |
| 8 | 0.201144 | 0.275103 |
| 9 | 0.190658 | 0.281981 |
| 10 | 0.221163 | 0.202201 |
| 11 | 0.238958 | 0.228336 |
| 12 | 0.289164 | 0.274415 |
| 13 | 0.350175 | 0.31293 |
| 14 | 0.485224 | 0.394085 |
| 15 | 0.566254 | 0.449794 |
| 16 | 0.705752 | 0.536451 |
| 17 | 0.835399 | 0.602476 |
| 18 | 0.811567 | 0.550894 |
| 19 | 0.763902 | 0.490371 |
| 20 | 0.629488 | 0.409904 |
| 21 | 0.580553 | 0.423659 |
| 22 | 0.705434 | 0.393398 |
| 23 | 0.716238 | 0.38033 |
| 24 | 0.530664 | 0.374828 |
| 25 | 0.47442 | 0.334938 |
| 26 | 0.515729 | 0.304677 |
| 27 | 0.506832 | 0.289546 |
| 28 | 0.408325 | 0.242779 |
| 29 | 0.387035 | 0.279917 |
| 30 | 0.462345 | 0.34663 |
| 31 | 0.466158 | 0.404402 |
| 32 | 0.385129 | 0.361073 |
| 33 | 0.409914 | 0.306052 |
| 34 | 0.472831 | 0.419532 |
| 35 | 0.468065 | 0.368638 |
| 36 | 0.489673 | 0.323934 |
| 37 | 0.330156 | 0.29436 |
| 38 | 0.352717 | 0.306052 |
| 39 | 0.30251 | 0.209078 |
| 40 | 0.224341 | 0.200138 |
| 41 | 0.204957 | 0.177442 |
| 42 | 0.227518 | 0.185695 |
| 43 | 0.254846 | 0.29436 |
| 44 | 0.303781 | 0.316369 |
| 45 | 0.387989 | 0.370014 |
| 46 | 0.546235 | 0.362448 |
| 47 | 0.498888 | 0.414718 |
| 48 | 0.45885 | 0.370014 |
| 49 | 0.441373 | 0.29436 |
| 50 | 0.35939 | 0.348693 |
| 51 | 0.338735 | 0.381018 |
| 52 | 0.333651 | 0.361761 |
| 53 | 0.385446 | 0.399587 |
| 54 | 0.391166 | 0.457359 |
| 55 | 0.435335 | 0.515131 |
| 56 | 0.382587 | 0.396149 |
| 57 | 0.344773 | 0.268226 |
| 58 | 0.392119 | 0.341128 |
| 59 | 0.398793 | 0.306052 |
| 60 | 0.35939 | 0.247593 |
| 61 | 0.370194 | 0.329436 |
| 62 | 0.3864 | 0.407153 |
| 63 | 0.377185 | 0.394085 |
| 64 | 0.358437 | 0.370702 |
| 65 | 0.364792 | 0.339752 |
| 66 | 0.408008 | 0.403026 |
| 67 | 0.411821 | 0.367263 |
| 68 | 0.401652 | 0.30055 |
| 69 | 0.643152 | 0.47868 |
| 70 | 0.693359 | 0.584594 |
| 71 | 0.488402 | 0.488308 |
| 72 | 0.342866 | 0.433975 |
| 73 | 0.285669 | 0.344567 |
| 74 | 0.225929 | 0.270977 |
| 75 | 0.203686 | 0.225585 |
| 76 | 0.205593 | 0.174003 |
| 77 | 0.21735 | 0.19326 |
| 78 | 0.268827 | 0.289546 |
| 79 | 0.306959 | 0.263411 |
| 80 | 0.363839 | 0.299862 |
| 81 | 0.363521 | 0.341816 |
| 82 | 0.428662 | 0.310179 |
| 83 | 0.423578 | 0.341816 |
| 84 | 0.452812 | 0.445667 |
| 85 | 0.426438 | 0.504814 |
| 86 | 0.364792 | 0.36795 |
| 87 | 0.376867 | 0.418845 |
| 88 | 0.405466 | 0.637552 |
| 89 | 0.358119 | 0.279917 |
| 90 | 0.324436 | 0.23934 |
| 91 | 0.373371 | 0.269601 |
| 92 | 0.380362 | 0.319807 |
| 93 | 0.328567 | 0.395461 |
| 94 | 0.340324 | 0.608666 |
| 95 | 0.385446 | 0.729711 |
| 96 | 0.376549 | 0.580468 |
| 97 | 0.364474 | 0.453232 |
| 98 | 0.276136 | 0.34663 |
| 99 | 0.260248 | 0.36795 |
| 100 | 0.255164 | 0.357634 |
| 101 | 0.238004 | 0.339065 |
| 102 | 0.20591 | 0.218019 |
| 103 | 0.240864 | 0.166437 |
| 104 | 0.25707 | 0.231774 |
| 105 | 0.345408 | 0.275103 |
| 106 | 0.372736 | 0.235213 |
| 107 | 0.389577 | 0.246905 |
| 108 | 0.40769 | 0.22077 |
| 109 | 0.430569 | 0.324622 |
| 110 | 0.45599 | 0.262724 |
| 111 | 0.46171 | 0.213205 |
| 112 | 0.455354 | 0.158872 |
| 113 | 0.390531 | 0.343191 |
| 114 | 0.455672 | 0.288858 |
| 115 | 0.487448 | 0.262036 |
| 116 | 0.370829 | 0.268913 |
| 117 | 0.363203 | 0.264787 |
| 118 | 0.449635 | 0.396836 |
| 119 | 0.397839 | 0.312242 |
| 120 | 0.34096 | 0.330124 |
| 121 | 0.408643 | 0.420908 |
| 122 | 0.446457 | 0.577717 |
| 123 | 0.387989 | 0.390646 |
| 124 | 0.397839 | 0.305365 |
| 125 | 0.476962 | 0.390646 |
| 126 | 0.469336 | 0.418845 |
| 127 | 0.418494 | 0.455296 |
| 128 | 0.448046 | 0.513067 |
| 129 | 0.367652 | 0.408528 |
| 130 | 0.354941 | 0.378955 |
| 131 | 0.292342 | 0.335626 |
| 132 | 0.239593 | 0.608666 |
| 133 | 0.228154 | 0.339752 |
| 134 | 0.230378 | 0.257909 |
| 135 | 0.292342 | 0.200138 |
| 136 | 0.504925 | 0.194635 |
| 137 | 0.485224 | 0.137552 |
| 138 | 0.932634 | 0.191197 |
| 139 | 1.01684 | 0.248281 |
| 140 | 0.698443 | 0.255846 |
| 141 | 0.611058 | 0.206327 |
| 142 | 0.514458 | 0.211829 |
| 143 | 0.387353 | 0.188446 |
| 144 | 0.337782 | 0.189821 |
| 145 | 0.347315 | 0.191197 |
| 146 | 0.334922 | 0.156809 |
| 147 | 0.376867 | 0.314305 |
| 148 | 0.35939 | 0.330812 |
| 149 | 0.312043 | 0.210454 |
| 150 | 0.34096 | 0.187758 |
| 151 | 0.378773 | 0.213893 |
| 152 | 0.358119 | 0.21458 |
| 153 | 0.359708 | 0.258597 |
| 154 | 0.418494 | 0.393398 |
| 155 | 0.454719 | 0.3989 |
| 156 | 0.435653 | 0.28542 |
| 157 | 0.447092 | 0.450481 |
| 158 | 0.411503 | 0.414718 |
| 159 | 0.328885 | 0.317056 |
| 160 | 0.330473 | 0.241403 |
| 161 | 0.2755 | 0.146492 |
| 162 | 0.24722 | 0.18707 |
| 163 | 0.283127 | 0.22696 |
| 164 | 0.316492 | 0.226272 |
| 165 | 0.40769 | 0.222146 |
| 166 | 0.459485 | 0.248281 |
| 167 | 0.484271 | 0.242091 |
| 168 | 0.529393 | 0.177442 |
| 169 | 0.468383 | 0.22696 |
| 170 | 0.480775 | 0.148556 |
| 171 | 0.366063 | 0.149931 |
| 172 | 0.351128 | 0.188446 |
| 173 | 0.408961 | 0.158184 |
| 174 | 0.390848 | 0.274415 |
| 175 | 0.335875 | 0.26685 |
| 176 | 0.384811 | 0.297111 |
| 177 | 0.438831 | 0.453232 |
| 178 | 0.400699 | 0.362448 |
| 179 | 0.345726 | 0.295736 |
| 180 | 0.392437 | 0.415406 |
| 181 | 0.483317 | 0.528198 |
| 182 | 0.47728 | 0.498624 |
| 183 | 0.453448 | 0.460798 |
| 184 | 0.538926 | 0.513755 |
| 185 | 0.486177 | 0.572215 |
| 186 | 0.351128 | 0.409904 |
| 187 | 0.31109 | 0.409216 |
| 188 | 0.288847 | 0.319807 |
| 189 | 0.274229 | 0.29436 |
| 190 | 0.306006 | 0.275791 |
| 191 | 0.34795 | 0.310867 |
| 192 | 0.391166 | 0.522696 |
| 193 | 0.324118 | 0.509629 |
| 194 | 0.47156 | 0.264099 |
| 195 | 0.435971 | 0.230399 |
| 196 | 0.424531 | 0.219395 |
| 197 | 0.462027 | 0.19945 |
| 198 | 0.536702 | 0.20564 |
| 199 | 0.518271 | 0.188446 |
| 200 | 0.483635 | 0.182256 |

|  | B-factor MD | B-Fact-xray |
| --- | --- | --- |
| 1 | 0.22891 | 0.323464 |
| 2 | 0.155636 | 0.293854 |
| 3 | 0.108875 | 0.256499 |
| 4 | 0.0684629 | 0.251334 |
| 5 | 0.0458756 | 0.207953 |
| 6 | 0.0450118 | 0.184369 |
| 7 | 0.0411675 | 0.205543 |
| 8 | 0.061529 | 0.238423 |
| 9 | 0.0966333 | 0.253916 |
| 10 | 0.151327 | 0.261319 |
| 11 | 0.287033 | 0.27664 |
| 12 | 0.469766 | 0.375452 |
| 13 | 0.698251 | 0.498365 |
| 14 | 0.592318 | 0.487175 |
| 15 | 0.49428 | 0.377518 |
| 16 | 0.370507 | 0.364262 |
| 17 | 0.227803 | 0.330177 |
| 18 | 0.239555 | 0.309348 |
| 19 | 0.228356 | 0.346015 |
| 20 | 0.246403 | 0.381821 |
| 21 | 0.173476 | 0.318471 |
| 22 | 0.126744 | 0.306593 |
| 23 | 0.154496 | 0.313651 |
| 24 | 0.15678 | 0.32398 |
| 25 | 0.0948422 | 0.29592 |
| 26 | 0.0857984 | 0.260802 |
| 27 | 0.135761 | 0.293854 |
| 28 | 0.109449 | 0.301429 |
| 29 | 0.0669566 | 0.268377 |
| 30 | 0.107542 | 0.27991 |
| 31 | 0.163966 | 0.293338 |
| 32 | 0.14006 | 0.300396 |
| 33 | 0.164905 | 0.282321 |
| 34 | 0.0928913 | 0.234119 |
| 35 | 0.106784 | 0.271475 |
| 36 | 0.0671065 | 0.256154 |
| 37 | 0.0338869 | 0.22293 |
| 38 | 0.0296587 | 0.246514 |
| 39 | 0.0452577 | 0.223963 |
| 40 | 0.161165 | 0.232226 |
| 41 | 0.308188 | 0.227922 |
| 42 | 0.126127 | 0.208814 |
| 43 | 0.243824 | 0.207609 |
| 44 | 0.200493 | 0.218282 |
| 45 | 0.131319 | 0.218798 |
| 46 | 0.140494 | 0.188673 |
| 47 | 0.187491 | 0.215355 |
| 48 | 0.209138 | 0.250646 |
| 49 | 0.172273 | 0.262007 |
| 50 | 0.241827 | 0.302978 |
| 51 | 0.249574 | 0.298674 |
| 52 | 0.258031 | 0.324841 |
| 53 | 0.196624 | 0.288518 |
| 54 | 0.154269 | 0.277673 |
| 55 | 0.172513 | 0.281116 |
| 56 | 0.199199 | 0.34722 |
| 57 | 0.143548 | 0.325701 |
| 58 | 0.121644 | 0.298158 |
| 59 | 0.184993 | 0.358409 |
| 60 | 0.184744 | 0.317783 |
| 61 | 0.125306 | 0.282665 |
| 62 | 0.154041 | 0.30229 |
| 63 | 0.234202 | 0.393011 |
| 64 | 0.19203 | 0.346359 |
| 65 | 0.145308 | 0.297642 |
| 66 | 0.133425 | 0.318471 |
| 67 | 0.083441 | 0.325701 |
| 68 | 0.0464976 | 0.266655 |
| 69 | 0.0326203 | 0.246686 |
| 70 | 0.0462483 | 0.23257 |
| 71 | 0.0677076 | 0.217766 |
| 72 | 0.0732388 | 0.23739 |
| 73 | 0.130481 | 0.244104 |
| 74 | 0.377588 | 0.252883 |
| 75 | 0.277194 | 0.271131 |
| 76 | 0.319211 | 0.289034 |
| 77 | 0.251893 | 0.311585 |
| 78 | 0.304663 | 0.341883 |
| 79 | 0.253348 | 0.306938 |
| 80 | 0.156551 | 0.27664 |
| 81 | 0.15885 | 0.31193 |
| 82 | 0.191777 | 0.344638 |
| 83 | 0.133848 | 0.306593 |
| 84 | 0.113507 | 0.316922 |
| 85 | 0.177112 | 0.345843 |
| 86 | 0.195087 | 0.382338 |
| 87 | 0.16964 | 0.364434 |
| 88 | 0.183254 | 0.413324 |
| 89 | 0.225873 | 0.485281 |
| 90 | 0.168688 | 0.357376 |
| 91 | 0.174442 | 0.398347 |
| 92 | 0.106406 | 0.296781 |
| 93 | 0.0930678 | 0.288174 |
| 94 | 0.0727696 | 0.278361 |
| 95 | 0.0476278 | 0.255121 |
| 96 | 0.0441562 | 0.240489 |
| 97 | 0.0668069 | 0.22775 |
| 98 | 0.118236 | 0.228955 |
| 99 | 0.132159 | 0.235152 |
| 100 | 0.134484 | 0.24221 |
| 101 | 0.183254 | 0.24135 |
| 102 | 0.149531 | 0.222241 |
| 103 | 0.118635 | 0.273369 |
| 104 | 0.151778 | 0.334137 |
| 105 | 0.293582 | 0.459115 |
| 106 | 0.289208 | 0.423997 |
| 107 | 0.275065 | 0.413841 |
| 108 | 0.248707 | 0.390429 |
| 109 | 0.273246 | 0.355655 |
| 110 | 0.247554 | 0.296092 |
| 111 | 0.18624 | 0.287657 |
| 112 | 0.205186 | 0.29592 |
| 113 | 0.222584 | 0.281632 |
| 114 | 0.162329 | 0.250818 |
| 115 | 0.134697 | 0.265278 |
| 116 | 0.128191 | 0.287141 |
| 117 | 0.147079 | 0.312618 |
| 118 | 0.129854 | 0.308315 |
| 119 | 0.100449 | 0.367189 |
| 120 | 0.118635 | 0.413152 |
| 121 | 0.139627 | 0.350146 |
| 122 | 0.106973 | 0.346531 |
| 123 | 0.154041 | 0.365123 |
| 124 | 0.108684 | 0.33276 |
| 125 | 0.10584 | 0.316406 |
| 126 | 0.0765657 | 0.256499 |
| 127 | 0.0699859 | 0.257876 |
| 128 | 0.101553 | 0.24376 |
| 129 | 0.116452 | 0.256843 |
| 130 | 0.199457 | 0.243071 |
| 131 | 0.218777 | 0.21811 |
| 132 | 0.289519 | 0.265967 |
| 133 | 0.295467 | 0.250473 |
| 134 | 0.344212 | 0.237218 |
| 135 | 0.474141 | 0.243071 |
| 136 | 0.48295 | 0.305388 |
| 137 | 0.375102 | 0.36168 |
| 138 | 0.368045 | 0.320881 |
| 139 | 0.320848 | 0.25581 |
| 140 | 0.427893 | 0.268205 |
| 141 | 0.417356 | 0.293682 |
| 142 | 0.411023 | 0.330177 |
| 143 | 0.323806 | 0.303495 |
| 144 | 0.249574 | 0.317611 |
| 145 | 0.250153 | 0.356688 |
| 146 | 0.284866 | 0.375968 |
| 147 | 0.271132 | 0.336891 |
| 148 | 0.231966 | 0.391978 |
| 149 | 0.139411 | 0.377518 |
| 150 | 0.149084 | 0.314684 |
| 151 | 0.209138 | 0.311241 |
| 152 | 0.15908 | 0.273025 |
| 153 | 0.151778 | 0.264589 |
| 154 | 0.15518 | 0.27664 |
| 155 | 0.174926 | 0.265622 |
| 156 | 0.265435 | 0.25822 |
| 157 | 0.35001 | 0.319504 |
| 158 | 0.342516 | 0.399552 |
| 159 | 0.389779 | 0.32157 |
| 160 | 0.339136 | 0.259253 |
| 161 | 0.384018 | 0.243071 |
| 162 | 0.397406 | 0.242383 |
| 163 | 0.332761 | 0.250646 |
| 164 | 0.32943 | 0.268721 |
| 165 | 0.392313 | 0.247891 |
| 166 | 0.377588 | 0.257531 |
| 167 | 0.328766 | 0.326734 |
| 168 | 0.420353 | 0.299707 |
| 169 | 0.476535 | 0.32157 |
| 170 | 0.398867 | 0.317955 |
| 171 | 0.393402 | 0.392839 |
| 172 | 0.350695 | 0.393872 |
| 173 | 0.264541 | 0.326218 |
| 174 | 0.223404 | 0.294543 |
| 175 | 0.238423 | 0.247375 |
| 176 | 0.172033 | 0.253228 |
| 177 | 0.194831 | 0.243243 |
| 178 | 0.212593 | 0.279566 |
| 179 | 0.257737 | 0.317611 |
| 180 | 0.201012 | 0.284903 |
| 181 | 0.297358 | 0.262352 |
| 182 | 0.27628 | 0.263557 |
| 183 | 0.244969 | 0.257531 |
| 184 | 0.290142 | 0.255466 |
| 185 | 0.365941 | 0.257531 |
| 186 | 0.29799 | 0.263729 |
| 187 | 0.310764 | 0.323464 |
| 188 | 0.412137 | 0.306765 |
| 189 | 0.435882 | 0.294371 |
| 190 | 0.377944 | 0.284042 |
| 191 | 0.303067 | 0.279738 |
| 192 | 0.239555 | 0.268205 |
| 193 | 0.181771 | 0.232226 |
| 194 | 0.157468 | 0.262868 |
| 195 | 0.111565 | 0.247719 |
| 196 | 0.109066 | 0.226889 |
| 197 | 0.0937755 | 0.244448 |
| 198 | 0.106029 | 0.211396 |
| 199 | 0.0817772 | 0.252883 |
| 200 | 0.0721463 | 0.247375 |

|  | B-Factor MD | B-Factor-xray |
| --- | --- | --- |
| 1 | 0.246691 | 0.883442 |
| 2 | 0.0887052 | 0.498802 |
| 3 | 0.0741818 | 0.44707 |
| 4 | 0.0638479 | 0.569216 |
| 5 | 0.0494129 | 0.461919 |
| 6 | 0.0374948 | 0.544308 |
| 7 | 0.039195 | 0.354942 |
| 8 | 0.0482616 | 0.303529 |
| 9 | 0.0705998 | 0.193358 |
| 10 | 0.0871603 | 0.258343 |
| 11 | 0.106029 | 0.359093 |
| 12 | 0.127157 | 0.33163 |
| 13 | 0.173717 | 0.154878 |
| 14 | 0.258031 | 0.353345 |
| 15 | 0.366992 | 0.350631 |
| 16 | 0.376522 | 0.441322 |
| 17 | 0.263054 | 0.361169 |
| 18 | 0.244969 | 0.216669 |
| 19 | 0.176625 | 0.332908 |
| 20 | 0.192538 | 0.330832 |
| 21 | 0.196367 | 0.319016 |
| 22 | 0.217695 | 0.343446 |
| 23 | 0.168925 | 0.411304 |
| 24 | 0.134484 | 0.363404 |
| 25 | 0.156093 | 0.350631 |
| 26 | 0.149979 | 0.324445 |
| 27 | 0.111759 | 0.315663 |
| 28 | 0.102663 | 0.372825 |
| 29 | 0.116254 | 0.467508 |
| 30 | 0.0993514 | 0.495449 |
| 31 | 0.0789872 | 0.468785 |
| 32 | 0.0907862 | 0.473894 |
| 33 | 0.099169 | 0.593326 |
| 34 | 0.0861379 | 0.434297 |
| 35 | 0.141799 | 0.450263 |
| 36 | 0.0645814 | 0.582947 |
| 37 | 0.0480076 | 0.438129 |
| 38 | 0.0477543 | 0.281814 |
| 39 | 0.040699 | 0.372665 |
| 40 | 0.034961 | 0.313588 |
| 41 | 0.0759263 | 0.266486 |
| 42 | 0.139844 | 0.316302 |
| 43 | 0.241258 | 0.295705 |
| 44 | 0.289831 | 0.414977 |
| 45 | 0.214466 | 0.222417 |
| 46 | 0.297043 | 0.199106 |
| 47 | 0.153361 | 0.265049 |
| 48 | 0.188244 | 0.200543 |
| 49 | 0.238706 | 0.313428 |
| 50 | 0.219861 | 0.298739 |
| 51 | 0.197909 | 0.376816 |
| 52 | 0.226148 | 0.400128 |
| 53 | 0.189754 | 0.543669 |
| 54 | 0.223678 | 0.632285 |
| 55 | 0.175896 | 0.349673 |
| 56 | 0.129646 | 0.338656 |
| 57 | 0.154496 | 0.403481 |
| 58 | 0.172033 | 0.469424 |
| 59 | 0.117839 | 0.288999 |
| 60 | 0.110024 | 0.393741 |
| 61 | 0.153361 | 0.35111 |
| 62 | 0.158619 | 0.328437 |
| 63 | 0.104901 | 0.426154 |
| 64 | 0.122859 | 0.505828 |
| 65 | 0.225598 | 0.512694 |
| 66 | 0.173959 | 0.596519 |
| 67 | 0.1309 | 0.714993 |
| 68 | 0.265733 | 0.87977 |
| 69 | 0.250153 | 0.779978 |
| 70 | 0.179066 | 0.519879 |
| 71 | 0.077691 | 0.499601 |
| 72 | 0.0570199 | 0.460003 |
| 73 | 0.0568817 | 0.378094 |
| 74 | 0.0461239 | 0.358295 |
| 75 | 0.0651712 | 0.411464 |
| 76 | 0.0712164 | 0.344244 |
| 77 | 0.0935984 | 0.48092 |
| 78 | 0.174442 | 0.492735 |
| 79 | 0.181277 | 0.449305 |
| 80 | 0.104526 | 0.488424 |
| 81 | 0.18925 | 0.50503 |
| 82 | 0.177599 | 0.435255 |
| 83 | 0.230574 | 0.288999 |
| 84 | 0.185242 | 0.453616 |
| 85 | 0.118037 | 0.401086 |
| 86 | 0.144647 | 0.483315 |
| 87 | 0.161863 | 0.416095 |
| 88 | 0.120838 | 0.483155 |
| 89 | 0.113507 | 0.397892 |
| 90 | 0.153587 | 0.354782 |
| 91 | 0.147523 | 0.294268 |
| 92 | 0.11764 | 0.439087 |
| 93 | 0.144207 | 0.54303 |
| 94 | 0.179556 | 0.608494 |
| 95 | 0.156322 | 0.521316 |
| 96 | 0.149979 | 0.418809 |
| 97 | 0.094486 | 0.434137 |
| 98 | 0.0780141 | 0.289638 |
| 99 | 0.066508 | 0.226409 |
| 100 | 0.0662097 | 0.330992 |
| 101 | 0.0509692 | 0.330353 |
| 102 | 0.0735525 | 0.311672 |
| 103 | 0.0698329 | 0.409229 |
| 104 | 0.0740242 | 0.449784 |
| 105 | 0.101553 | 0.440045 |
| 106 | 0.195342 | 0.450263 |
| 107 | 0.154041 | 0.361328 |
| 108 | 0.129646 | 0.377136 |
| 109 | 0.162796 | 0.489222 |
| 110 | 0.126744 | 0.435255 |
| 111 | 0.14006 | 0.367396 |
| 112 | 0.137902 | 0.353185 |
| 113 | 0.149531 | 0.510299 |
| 114 | 0.110601 | 0.399808 |
| 115 | 0.104713 | 0.357496 |
| 116 | 0.13258 | 0.494491 |
| 117 | 0.137687 | 0.477088 |
| 118 | 0.116254 | 0.434456 |
| 119 | 0.11234 | 0.507425 |
| 120 | 0.152003 | 0.505349 |
| 121 | 0.161398 | 0.407472 |
| 122 | 0.121846 | 0.467508 |
| 123 | 0.138764 | 0.466071 |
| 124 | 0.188495 | 0.622226 |
| 125 | 0.182265 | 0.49034 |
| 126 | 0.146635 | 0.555644 |
| 127 | 0.174442 | 0.608973 |
| 128 | 0.260094 | 0.622226 |
| 129 | 0.120838 | 0.39406 |
| 130 | 0.0932445 | 0.363723 |
| 131 | 0.0741818 | 0.3848 |
| 132 | 0.0529484 | 0.357337 |
| 133 | 0.054154 | 0.372505 |
| 134 | 0.0548296 | 0.458407 |
| 135 | 0.0837757 | 0.594284 |
| 136 | 0.121443 | 0.508702 |
| 137 | 0.299572 | 0.330034 |
| 138 | 0.252475 | 0.483155 |
| 139 | 0.267227 | 0.521316 |
| 140 | 0.294838 | 0.5384 |
| 141 | 0.401063 | 0.569855 |
| 142 | 0.743962 | 0.486348 |
| 143 | 0.362796 | 0.636277 |
| 144 | 0.259799 | 0.633882 |
| 145 | 0.324136 | 0.57704 |
| 146 | 0.372978 | 0.60099 |
| 147 | 0.384018 | 0.777104 |
| 148 | 0.234202 | 0.62494 |
| 149 | 0.182759 | 0.612805 |
| 150 | 0.252766 | 0.594923 |
| 151 | 0.252184 | 0.4616 |
| 152 | 0.173959 | 0.549098 |
| 153 | 0.163732 | 0.52882 |
| 154 | 0.191523 | 0.584863 |
| 155 | 0.160933 | 0.473735 |
| 156 | 0.176138 | 0.510618 |
| 157 | 0.133848 | 0.521954 |
| 158 | 0.0986229 | 0.310075 |
| 159 | 0.100266 | 0.476449 |
| 160 | 0.065319 | 0.460163 |
| 161 | 0.060528 | 0.393741 |
| 162 | 0.0555095 | 0.445314 |
| 163 | 0.110216 | 0.432221 |
| 164 | 0.168212 | 0.380808 |
| 165 | 0.264243 | 0.382405 |
| 166 | 0.445489 | 0.368354 |
| 167 | 0.442788 | 0.433977 |
| 168 | 0.485367 | 0.436532 |
| 169 | 0.591873 | 0.376976 |
| 170 | 0.434736 | 0.416574 |
| 171 | 0.33176 | 0.464793 |
| 172 | 0.479737 | 0.425355 |
| 173 | 0.529479 | 0.471659 |
| 174 | 0.356201 | 0.416733 |
| 175 | 0.385454 | 0.453137 |
| 176 | 0.593655 | 0.539358 |
| 177 | 0.540913 | 0.632285 |
| 178 | 0.390141 | 0.532492 |
| 179 | 0.369099 | 0.47597 |
| 180 | 0.284866 | 0.4517 |
| 181 | 0.148637 | 0.368034 |
| 182 | 0.0984412 | 0.424397 |
| 183 | 0.0849527 | 0.313588 |
| 184 | 0.0582707 | 0.450902 |
| 185 | 0.0637017 | 0.463197 |
| 186 | 0.0875024 | 0.264091 |
| 187 | 0.136402 | 0.409229 |
| 188 | 0.166082 | 0.347916 |
| 189 | 0.248419 | 0.351269 |
| 190 | 0.237858 | 0.253872 |
| 191 | 0.136188 | 0.372505 |
| 192 | 0.170356 | 0.328916 |
| 193 | 0.216347 | 0.362925 |
| 194 | 0.15885 | 0.410346 |
| 195 | 0.148414 | 0.40875 |
| 196 | 0.231966 | 0.383363 |
| 197 | 0.23673 | 0.373942 |
| 198 | 0.216077 | 0.422801 |
| 199 | 0.122453 | 0.467667 |
| 200 | 0.123875 | 0.429028 |

**Figure SF6.** B-factor comparison between MD and X-ray (**A**) 3E96 (inactive) (**B**) YAGE (**C**) KDGA (**D**) NAL

**Major structural motions**

Principle component Analysis (PCA) a multivariate technique based essential dynamics was used for extracting large scale concerted motions from the MD trajectories, of proteins belong to 3E96 (inactive), YAGE, KDGA and NAL families.

**Motions in 3E96**

Motion vectors of segments of interest, Loop A and Loop B, drift away from each other towards the periphery. Likewise Loop B1 drifts away from Loop A. These motions further open the IS1 region which is already wide open in the crystal structure. Motion vectors of Loop A2 indicates that this segment moves closer to loop B1 and loop B. This indicates that IS2 region gets further tightened compared to the crystal structure.

**Figure SF7:** **(A)** Major motions in protein 3E96 captured by PCA analysis showing loop A, loop B and loop B1 moves towards periphery there by drifting away from the core. The motion vectors of loop A2 pointing upwards indicate that this segments moves closer to loop B1 and loop B. **(B)** Major motions of segments in interest is shown in schematic representation for clarity.

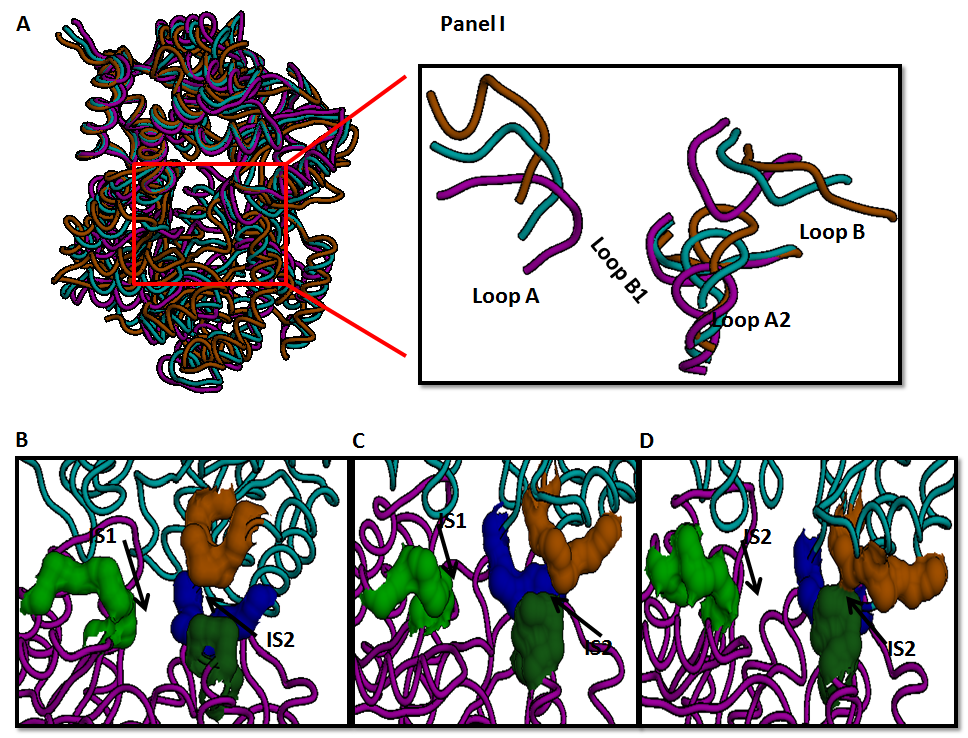

**Figure SF8:** Comparison of 0^th^, 49^th^ and 99^th^ (give in terms of time ps) frames of molecular dynamics simulations of protein 3E96 **(A)** Image showing superimposition of 0^th^ (Magenta), 49^th^ (Cyan) and 99^th^ frames (Orange). Image shown in inset that focus the segments under analysis clearly shows loop A and loop B drifts away from each other as simulation progress from 0^th^ to 99^th^ frame. Image also shows that loop A2 moves closer to loop B1 and loop B. **(B)** Image showing surfaces of Loop A (Green), loop B (Orange), loop B1 (Blue) and loop A2 (Dark green) in 0^th^ frame. IS1 and IS2 regions in core and periphery respectively are indicated by arrows. **(C)** Image showing surfaces of the segments in 49^th^ frame. **(D)** Images showing surfaces of the segments in 99^th^ frame. The widening of gap between the surfaces of loop A (green) and loop B (orange) as simulation progresses depicts the drifting of the segments from core towards the periphery, thereby opening the IS1 region. The closing up of the surfaces of loop A2 (dark green) and loop B (orange) as simulation progresses indicates that IS2 region is getting tightened.

**Motions in KDGA protein (PDBid-1W37)**

Motion vectors of segments of interest, Helix A and Helix B makes an initial motion towards the core and moves away towards the periphery. Thus they exhibit sliding motion with respect to each other. The sliding motion brought them closer initially and then allowed them to drift away. In addition, the initial motion towards the core brought Helix A closer to the motionless loop B1. The initial close approach of helix A and B and the close approach of loop A with loop B1 maintain the tightness of IS1 region. Though loop B1 is relatively motionless, the motion vectors seen in the extreme right end of this segment move towards periphery in parallel with helix A2, therefore the two segments did not come close or drift away. Also the motion of helix B towards the periphery, brought the segment closer to helix A2, however the decrease in distance was not sufficient to make surface contact. Thus IS2 region is largely unperturbed throughout the simulation.

**Figure SF9: (A)** Major motions in protein 1W37 captured by PCA analysis showing Helix A and Helix B making an initial movement towards the core and later move towards periphery. Motion vector indicates that loop B1 is largely static except for the few motion vectors seen in the extreme right end towards periphery. Motion vectors on helix A2 indicate that they move towards periphery **(B)** Schematic representation of the motion vectors is shown for clarity.

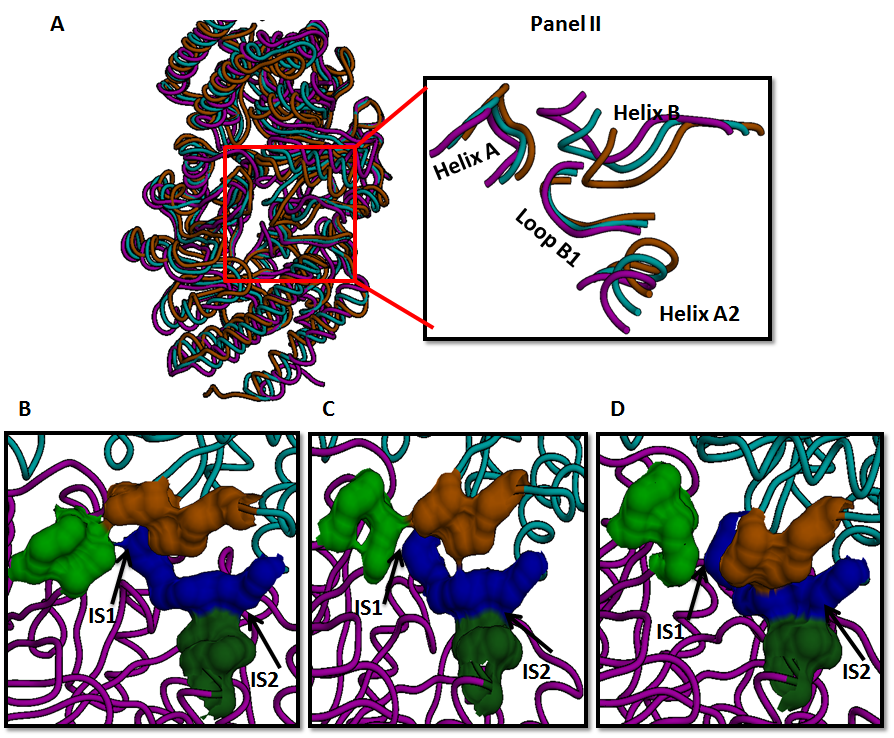

**Figure SF10:** Comparison of 0^th^, 49^th^ and 99^th^ frames (give in terms of time ps) of molecular dynamics simulations of protein 1W37 **(A)** Image showing superimposition of 0^th^ (Magenta), 49^th^ (Cyan) and 99^th^ frames (Orange). Image shown in inset focuses the segments under analysis **(B)** Image showing surfaces of Loop A (Green), loop B (Orange), loop B1 (Blue) and loop A2 (Dark green) in 0^th^ frame. IS1 and IS2 regions in core and periphery respectively are indicated by arrows. **(C)** Image showing surfaces of the segments in 49^th^ frame. **(D)** Images showing surfaces of the segments in 99^th^ frame. Initial movement of helix A and helix B, results in closing up of the segments, as is evident from the undisturbed surface contact between the 2 segments in the 49^th^ frame. Later movement of drifting towards the periphery is evident from the ruining of the surface contact as well as the widening gap between the 2 segments could be seen in 99^th^ frame. Helix A and B exhibiting a sliding motion with respect to each other is evident from the change in scenario in 0^th^ frame, wherein the surfaces of helix A (green) and helix B (orange) shown juxtaposed moved, with the former moving upwards and the later taking a downward motion in 49^th^ and 99^th^ Frame. The initial close up of the segments and static loop B1 maintained or slightly tightened the IS1 region. Helix A2 taking an upward motion towards periphery though brought it closer to loop B1, the concave trough is maintained throughout the simulation. The upward and downward motion towards periphery undertook by helix A2 (dark green) and helix B (orange) respectively, though brought them closer, their surfaces do not come in contact, thereby leaving the IS2 unperturbed as seen in 99^th^ frame

**Motions in NAL protein (PDBid-1NAL)**

Motion vectors on loop B and loop B1 indicates that they move towards core and remain close to loop A, which itself stays static in the core except for a mild twisting motion as depicted by the clockwise movement of arrows on loop A. The movement of loop B and loop B1 towards core and loop A remaining in core tighten the IS1 region as compared to crystal structure. Motion vectors on loop A2 shows that the segment moves towards the periphery, however the movement of loop B1 towards the core maintains the distance between the segments or brings them slightly closer as compared to crystal structure. A small closing of the segments is not sufficient to make any significant increase in the already negligible IS2 region seen in crystal structure.

**Figure SF11: (A)** Major motions in protein 1NAL captured by PCA analysis showing loop A, loop B and loop B1 are largely static. The motion of right end of the loop B and loop B1 indicate that they move towards the core while the mild twisting of loop A indicates that the segments do not drift but stays in its original position. The motion vectors of loop A2 indicates that it move towards periphery. **(B)** Schematic representation of the motion vectors is shown for clarity.

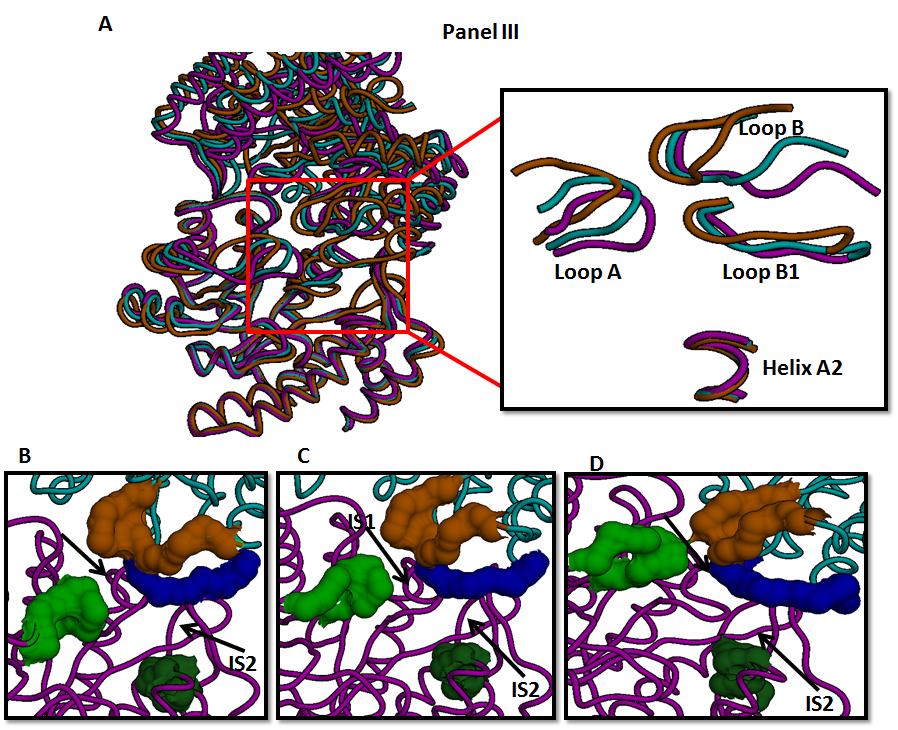

**Figure SF12:** Comparison of 0^th^, 49^th^ and 99^th^ frames (give in terms of time ps) of molecular dynamics simulations of protein 1NAL **(A)** Image showing superimposition of 0^th^ (Magenta), 49^th^ (Cyan) and 99^th^ frames (Orange). Image shown in inset focuses the segments under analysis **(B)** Image showing surfaces of Loop A (Green), loop B (Orange), loop B1 (Blue) and loop A2 (Dark green) in 0^th^ frame. IS1 and IS2 regions in core and periphery respectively are indicated by arrows. **(C)** Image showing surfaces of the segments in 49^th^ frame. **(D)** Images showing surfaces of the segments in 99^th^ frame. The twisting motion at the right and left end of the loop A and the movement of extreme right end of the loop B in the ‘open the lid’ fashion push the interfacing middle portion of the segments closer to each other, which is evident from the narrowing of the gap between the segments as simulation progresses that culminates in surface contact in the 99^th^ frame. The twisting motion of loop A that pushes the interfacing portion of that segment towards the core brought it closer to largely motionless loop B1. Thus loop A moves closer to loop B and loop B1 resulting in tightening of IS1 as compared to crystal structure. Motions of loop A2 and extreme right end of loop B1, which are antiparallel to each other maintains a constant distance between the two segments and thus they do not close up to form IS2 region throughout the simulation period. The increase in gap between loop B (orange) and loop B1 (blue) clearly depicts the movement of right end of loop B1 towards the core in a ‘open the lid’ fashion, which also results in increased distance from loop A2.

**Motion in protein 2V8Z (a KDGA homolog)**

Motion vectors on helix A and helix B indicates they undergo twisting motion with respect to each other. Twisting motion keeps the two segments from drifting away and to stay in core. The twisting motion of helix A, makes it move away from the motionless loop B1 and may weaken the IS1 region interactions, the closing of helix A and B due to twisting motion maintains the tight interactions of IS1. The relatively motionless loop B1 and helix A2 maintain the distance between the two segments, thereby leaving the IS2 region unperturbed.

**Figure SF13: (A)** Major motions in protein 2V8Z captured by PCA analysis showing Helix A and Helix B undergo twisting motion with respect to each other. This indicates that the segments do not drift away but remain in the core. Absence of any motion vectors on loop B1 and helix A2 indicates the segments are largely static. **(B)** Schematic representation of the motion vectors is shown for clarity.

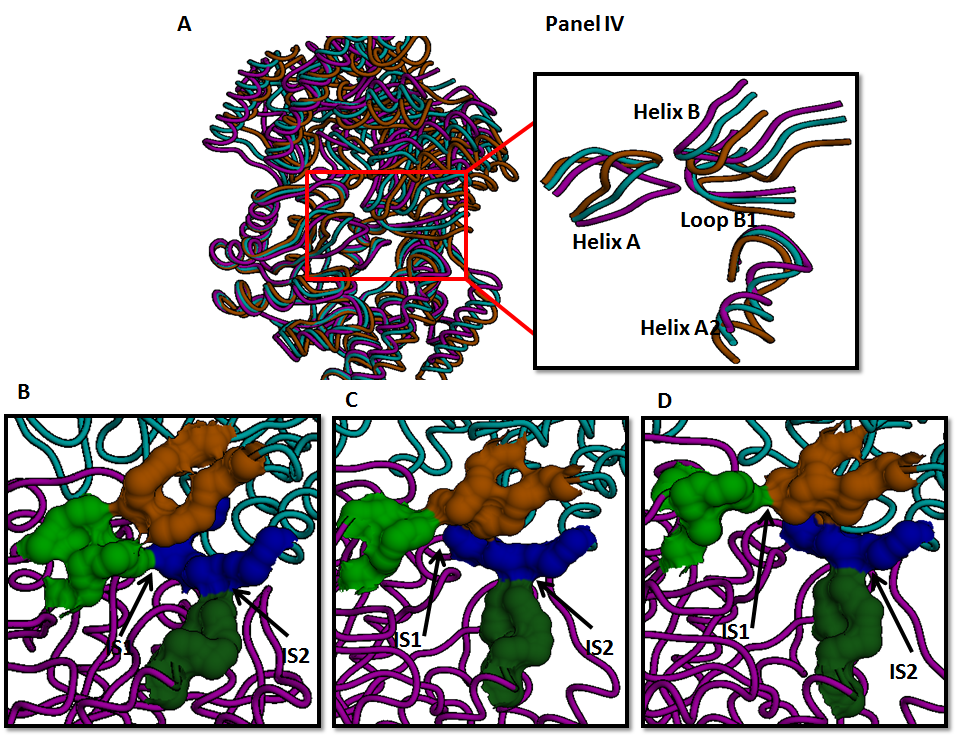

**Figure SF14:** Comparison of 0^th^, 49^th^ and 99^th^ frames (give in terms of time ps) of molecular dynamics simulations of protein 2V8Z **(A)** Image showing superimposition of 0^th^ (Magenta), 49^th^ (Cyan) and 99^th^ frames (Orange). Image shown in inset focuses the segments under analysis **(B)** Image showing surfaces of Loop A (Green), loop B (Orange), loop B1 (Blue) and loop A2 (Dark green) in 0^th^ frame. IS1 and IS2 regions in core and periphery respectively are indicated by arrows. **(C)** Image showing surfaces of the segments in 49^th^ frame. **(D)** Images showing surfaces of the segments in 99^th^ frame. Loss of surface contact between left end of loop B1 and helix A in 49^th^ and 99^th^ frame is due to widening distance between the segments due to twisting motion of helix A. The twisting motion of helix A and B with respect to each other holds the segments from drifting away, which is evident from the surface contact being maintained in 99^th^ frame. Though the twisting motion of helix A weakens the IS1 it shares with loop B1, the surface contact maintained with helix B ensures the steric crowding of IS1. The lack of motion of loop B1 and helix A2 leave the IS2 region unperturbed throughout the simulation period.

**Cavity volume analysis in MD simulation:**

The change in the cavity volume during simulation time was monitored using EPOCK. The variation in the active site for the four proteins (3E96, KDGA, NAL and YAGE) was calculated in a time dependent manner using epock program. The active site was defined from the center of the pocket picking CG2 atom of Ile116 in 3E96, OG1 atom of Thr44 in 1W37 (KDGA), CE1 atom of Phe252 in 1NAL (NAL) and CE2 atom of Phe265 in 2V8Z (YagE). The grid spacing was set at 0.5 Angstrom. A box size of 30 ×16 ×14 is defined for 3E96 and a box size of 14×14×14 is defined for the rest of the proteins.

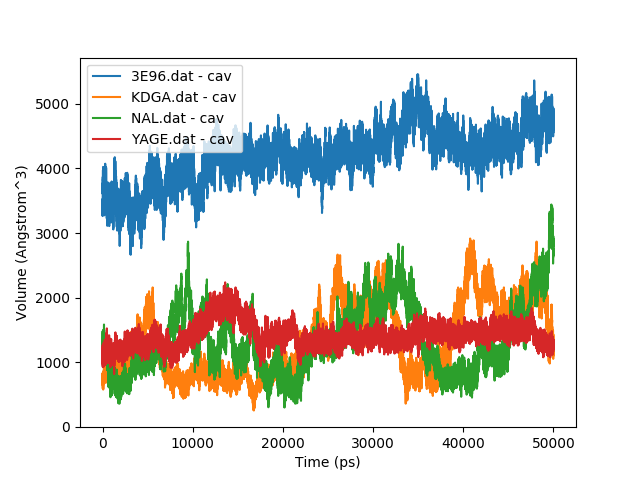

**Figure SF15:** Plot showing change in cavity volume with respect to time. 3E96 (blue), KDGA (Orange), NAL (Green); YagE (Red)
